# Probing remdesivir nucleotide analogue insertion to SARS-CoV-2 RNA dependent RNA polymerase in viral replication

**DOI:** 10.1101/2021.07.12.452099

**Authors:** Moises Ernesto Romero, Chunhong Long, Daniel La Rocco, Anusha Mysore Keerthi, Dajun Xu, Jin Yu

## Abstract

Remdesivir (RDV) prodrug can be metabolized into a triphosphate form nucleotide analogue (RDV-TP) to bind and insert into the active site of viral RNA dependent RNA polymerase (RdRp) to further interfere with the viral genome replication. In this work, we computationally studied how RDV-TP binds and inserts to the SARS-CoV-2 RdRp active site, in comparison with natural nucleotide substrate adenosine triphosphate (ATP). To do that, we first constructed atomic structural models of an initial binding complex (active site open) and a substrate insertion complex (active site closed), based on high-resolution cryo-EM structures determined recently for SARS-CoV-2 RdRp or non-structural protein (nsp) 12, in complex with accessory protein factors nsp7 and nsp8. By conducting all-atom molecular dynamics simulation with umbrella sampling strategies on the nucleotide insertion between the open and closed state RdRp complexes, our studies show that RDV-TP can bind comparatively stabilized to the viral RdRp active site, as it primarily forms base stacking with the template Uracil nucleotide (at +1), which is under freely fluctuations and supports a low free energy barrier of the RDV-TP insertion (∼ 1.5 kcal/mol). In comparison, the barrier (*∼* 2.6 kcal/mol), when the fluctuations of the template nt are well quenched. The simulations also show that the initial base stacking of RDV-TP with the template can be particularly stabilized by motif B-N691, S682, and motif F-K500 with the sugar, base, and the template backbone, respectively. Although the RDV-TP insertion can be hindered by motif-F R555/R553 interaction with the triphosphate, the ATP insertion seems to be facilitated by such interactions. The inserted RDV-TP and ATP can be further distinguished by specific sugar interaction with motif B-T687 and motif-A D623, respectively.

## 2 Introduction

RNA dependent RNA polymerase (RdRp) is the core protein engine responsible for synthesizing genome in the replication/transcription machinery of RNA viruses, which represent a large class of human and animal pathogens to cause disease and pandemics [1, 2]. Based on the template RNA strand, RdRp selectively recruits ribonucleotides one at a time to the active site and adds the nucleotide to the growing RNA chain upon catalyzing the phosphoryl-transfer reaction, which is then followed by product (pyrophosphate) release and the polymerase translocation. Due to its critical role in the viral RNA synthesis and highly conserved core structure, the viral RdRp serves a highly promising antiviral drug target for both nucleotide analogue and non-nucleoside inhibitors [3]. Remdesivir (or RDV), the only FDA proved drug (named VEKLURY) so far treating COVID-19 [4], works as a prodrug that is metabolized into a nucleotide analogue to compete with natural nucleotide substrates of RdRp to be incorporated into viral RNA gnome to further terminate the RNA synthesis [5,6]. As a broad-spectrum anti-viral compound, RDV was developed originally for treatments of Ebola virus disease (EVD) [7], and then applied for infections of middle east and severe accurate respiratory syndrome coronavirus (MERS-CoV and SARS-CoV) [8], which are both close relatives to the currently emerged novel coronavirus (SARS-CoV-2) causing COVID-19. Recent in-vitro and in-vivo studies on RDV impacts to the viral RdRp function have confirmed the RDV analogue incorporation during the viral RdRp replication, in particular, in SARS-CoV-2 [9–11]. The existing evidences have consistently suggested that the active triphosphate form of RDV (RDV-TP) binds competitively with the natural substrate, i.e., adenosine triphosphate (or ATP), to the viral RdRp and the incorporation leads to a delayed chain termination [9]. Such an analogue incorporation and consequent chain termination indicate that RDV-TP can successfully evade from both nucleotide selectivity of the viral RdRp as well as the proofreading function from ExoN in coordination with RdRp in the coronavirus replication [5,12].

Nucleotide selectivity of the RdRp or polymerases in general serves as a primary fidelity control method in corresponding gene transcription or replication, i.e., during the template-based polymerase elongation [13–15]. The selectivity indeed proceeds throughout a full nucleotide addition cycle (NAC), consisting of nucleotide substrate initial binding, insertion to the active site, catalysis, product (or pyrophosphate) release, and together with the polymerase translocation [16]. To be successfully incorporated, the antiviral nucleotide analogue needs to pass almost every fidelity checkpoint in the polymerase NAC [17, 18]. In coronaviruses with large genome sizes, proofreading conducted by an exonuclease (or ExoN) protein further improves the RNA synthesis fidelity [12]. Correspondingly, the nucleotide analogue drug need further evade from the ExoN proofreading to ultimately terminate the RdRp elongation. Although RDV succeeds as a nucleotide analogue drug to interfere with the CoV-2 RdRp function, as being demonstrated in vivo and in vitro, the underlying structural dynamics mechanisms on how that being achieved are still to be determined, and *in silio* approaches may particularly help. Recent modeling and computational efforts have been made to approach the underlying mechanisms of the RDV-TP binding and incorporation to the CoV-2 RdRp, from molecular docking [19] and binding free energy calculation upon the nucleotide initial association [20], to nucleotide addition together with potential ExoN proofreading activities [21]. Nevertheless, those studied structural systems were still made by constructing homology model of the SARS-CoV-2 RdRp according to a previously resolved structure of the SARS-CoV RdRp [22]. Upon very recent high-resolution cryo-EM structures being resolved on the SARS-CoV2 RdRp (the non-structural protein or nsp12), with and without incorporation of RDV [23,24], it becomes highly desirable to conduct all-atom modeling and molecular dynamics (MD) simulations directly on the CoV-2 RdRp structure, so that to probe how RDV succeeds at binding and inserting into the RdRp active site, despite of existing nucleotide selectivity of RdRp to be against non-cognate nucleotide species [15].

The high-resolution structures of SARS-CoV-2 RdRp or nsp12 were obtained in complex with accessory protein nsp7 and nsp8, which are supposed to assist processivity of the replication/transcription machinery along the viral RNA [25] (see **Fig 1A**). The core RdRp (residue 367-920, excluding the N-terminal NiRAN and interfacial region) adopts a handlike structure, consisting of fingers, palm, and thumb subdomains, similar to other single-subunit viral RNA polymerases (RNAPs) and family-A DNA polymerases (DNAPs) [26–28]. There are seven highly conserved structural motifs shared by RdRps, located in the palm (A-E) and fingers (F-G) subdomains (**Fig 1B**). In general, when there is no substrate bound, the RdRp active site adopts an open conformation. A nucleotide substrate can bind to the active site in the open conformation, and inserts into the active site to reach a closed conformation, as the nucleotide is stabilized or to be ready for the catalytic reaction [29,30]. In the recently resolved SARS-CoV-2 RdRp structures, both the open and closed conformation state of the active site were captured, with the former in the absence of the substrate [24], and the latter captured with an RDV analogue already incorporated to the end of the synthesizing RNA chain (i.e., in post-catalytic or product state) [23]. In order to probe how a nucleotide or analogue binds and inserts to the RdRp active site, we accordingly constructed both an open (i.e. substrate initial binding) and a closed (substrate insertion) structural complex of the CoV-2 RdRp, based on the newly resolved structures (PDB: 7BTF [24] and 7BV2 [23]) (see **Fig 1C** for a closed form). Note that in the single-subunit viral RNAPs or DNAPs, the nucleotide insertion, in accompany with the open to closed conformational transition (or pre-chemistry transition or isomerization), usually happens slowly (e.g. milliseconds or above), i.e., to be rate limiting (or partially rate-limiting) in the NAC [31–33]. Such a slow nucleotide insertion step correspondingly plays a significant role in the nucleotide selection or fidelity control, for example, in the single-subunit viral T7 RNAP system studied recently [34–36]. To understand how RDV-TP can evade from the nucleotide selectivity of RdRp to be incorporated, it is therefore essential to probe how such a nucleotide analogue binds stably and inserts sufficiently fast or with low energy barrier into the active site, comparing to its natural substrate counterpart. Accordingly, in this work, we employed all-atom MD simulation to probe mainly the free energetics of the RDV-TP insertion into the CoV-2-RdRp active site, in comparison with the ATP insertion. To do that, umbrella sampling strategies were implemented connecting the initial substrate binding (active site open) and the insertion (active site closed) conformational states, in particular, by enforcing collective coordinates of atoms from structural motif A-G and the inserting NTP (excluding or including the template nucleotide or nt +1 with forcing), i.e., from the active site open to the closed state. The simulations consequently reveal free energetics or potentials of mean force (PMFs) along the reaction coordinate of the RDV-TP and ATP insertion, demonstrating how local residues around the RdRp active site or NTP binding site coordinate with the nucleotide binding insertion and differentiation, comparing RDV-TP and ATP.

**Figure 1:**
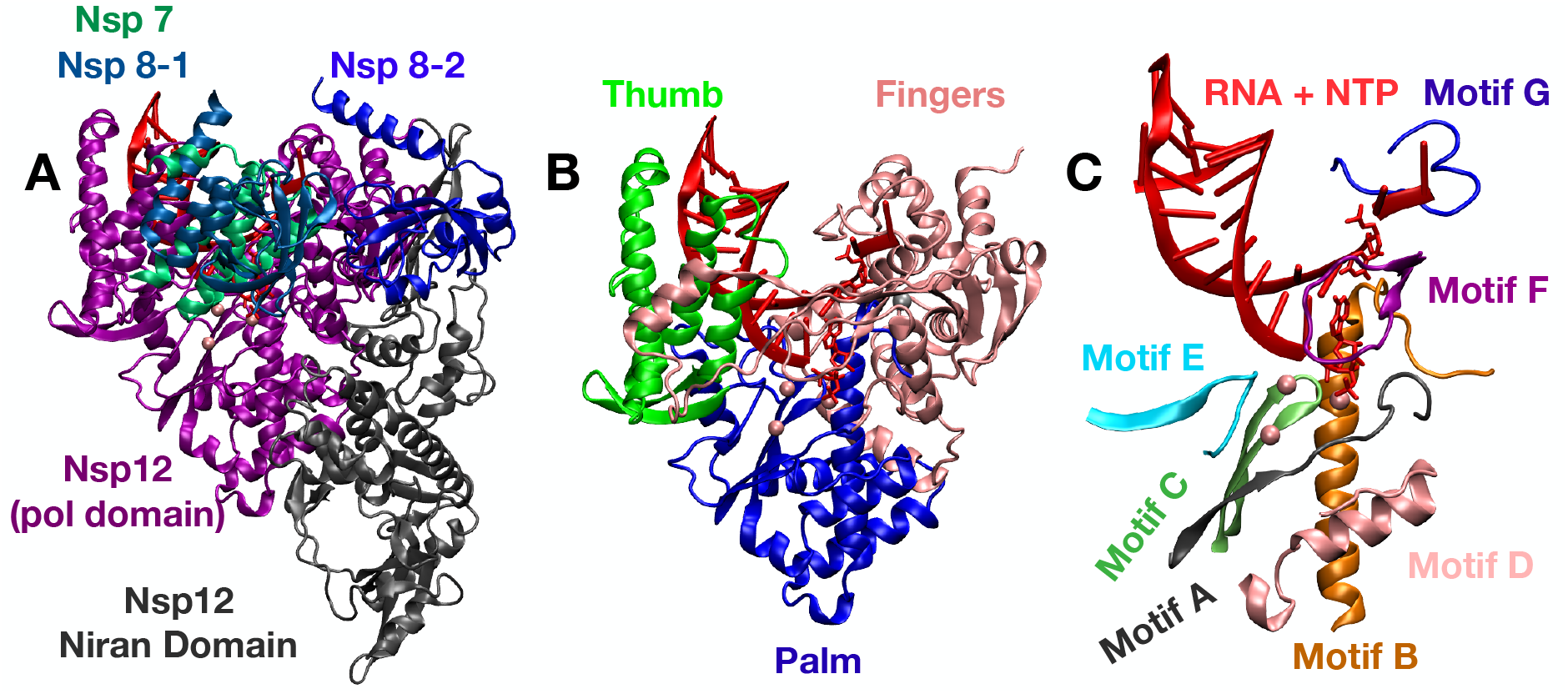
SARS-CoV-2 RdRp elongation complex with an incorporated remdesivir (RDV) in the closed state (baed on PDB:7BV2 [23]). **A** The two main domains (N-terminus domain in grey, polymerase or pol domain in purple) of the RdRp along with the three cofactors (nsp8’s in blue and nsp7 in green). **B** The pol domain consists three subdomains thumb (green), fingers (pink), and palm (blue). RNA (red) and NTP +1 Template (red licorice). **C** Motifs A-G within the pol domain.

## 3 Computational Details

### 3.1 Building Open/Closed structures and RDV Force Field

High-resolution Cryo-EM structures for CoV-2-RdRp’s elongation complex are available in a post-catalysis state with RDV analogue incorporated (PDB:7BV2) [23]. Using this structure, RDV-TP and ATP were fitted into the active site to create the closed /substrate binding complex (see **Supplementary Information** or **SI Methods** for details). Missing residues were added using MODELLER 9.24 [37] and an apo nsp12 structure as a reference (PDB:7BTF) [24] The open state was constructed from the apo nsp12 structure [24] along with additionally incorporated RNA strands and RDV-TP (or ATP) by fitting the above constructed RdRP closed structure with the apo RdRp structure.

Histidine protonation states were predicted using PDB2PQR [38] and PROPKA3 [39] followed by visual inspection. The two nsp8 N-terminals were cleaved and shorted by 11 residues (see **SI Methods**). A force field was generated for RDV, with partial charges calculated by following the formalism used in amber nucleic acid forcefields [40]. RDV-TP 3’ and 5’ terminals were truncated, and replaced with terminal hydroxyl groups (see **SI Methods** and **Fig S1**). A Hartree-Fock calculation at the level of HF/6-31G* was set to perform geometric optimization and a self-consistent calculation to obtain an electro-static potential for constrained charge fitting [41]. Using the two-stage Restrained Electrostatic Potential method [42], partial atomic charges for the RDV were generated(see **SI Table S1** for more details). Torsional parameters were taken from Parmbsc1 when applicable and the general Amber force field (GAFF) [43]. RDV force field parameters were constructed using antechamber [44]. Adaptive Poisson-Boltzmann Solver (APBS) mapping of the modeled substrate structures are provided in **SI Fig S2**. Docking of RDV-TP and ATP has also been conducted to the open state RdRp complex to compare and confirm with the constructed initial binding structural complex of RDV-TP or ATP(see **SI Methods Fig S3-4**).

### 3.2 Simulation Details

All MD simulations were performed using Gromacs 2019 package [45] with the Amber14sb protein force field [46] and Parmbsc1 nucleic acid parameters [47]. For the NTPs, triphosphate parameters calculated previously were used [48]. Each of the RdRp complexes was solvated with explicit TIP3P water [49] with a minimum distance from the complex to the wall set to 15Å, resulting in an average box size of 15.7nm x 15.7 nm x 15.7 nm (see **SI Fig S5**). Sodium and chloride ions were added to neutralize the systems and make the salt concentration 100mM. Three magnesium ions were kept from the cryo-EM structures [23]. The full simulation systems contained on average about 382,000 atoms. For all simulations, the cut-off of van der Waals (vdw) and the short range electrostatic interactions were set to 10Å. Particle-mesh-Ewald (PME) method [50, 51] was used to evaluate the long-range electrostatic interactions. Timestep was 2fs and the neighbor list was updated every 10 steps. Temperature was kept at 310 K using the velocity re-scaling thermostat [52]. Pressure was kept at 1 bar using Berendsen barostat [53] during pre-equilibration and Parrinello-Rahman barostat [54] for production, targeted MD (TMD), and umbrella simulation runs. Each initial system was minimized for a maximum of 50000 steps using steepest-descent algorithm, followed by a 2-ns NVT MD simulation with all the heavy atoms in the system fully constrained. Next a 2-ns NPT simulation along with the same constraints was performed. Constraints were released in 1-ns intervals in the following order: RNA, nsp8/nsp-7, nsp12/NTP/metal ions. Equilibration is then conducted for 100 ns or longer. TMD and umbrella sampling simulations were accomplished by using gromacs patched with plumed 2.6.1 [55]

### 3.3 Determining the Reaction Coordinate and Calculating Free Energy

The open to close conformational change of the RdRp is expected to be on the order of milliseconds and therefore can not be captured by brute force MD. In order to calculate free energy, the umbrella sampling method was used. [56–58] To use such a method a reaction path needs to be provided. In this study we used TMD to generate such a path between the open and closed states. TMD [59] implementation requires an initial and a final structures to be specified which we continue using in the umbrella sampling simulations. In this work we implemented two slightly varied protocols by manipulating coordinates of two slightly varied atom sets: nsp12 motifs (motif A-G backbone atoms) and NTP (heavy atoms) or nsp12 motifs (motif A-G backbone atoms), NTP (heavy atoms), and template +1 nucleotide (heavy atoms). The corresponding RC is then constructed by aligning the structures to reference structures via the fingers sub-domain and measuring the differences of RMSDs.

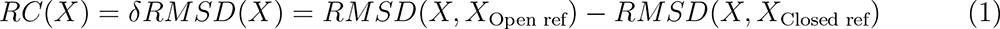

Where X is the coordinates for the above selected atom sets and *X*_Open ref_/*X*_Closed ref_ is a chosen reference state (see **SI Methods**).

#### 3.3.1 Target MD and Umbrella Sampling

Using the selected open and closed reference states TMD is launched from each state to create paths (forward path started from the open to the closed reference structure, and the backward path started from the closed then to the open reference structure) that meet halfway on the RC (see **SI Fig S6** and **movies S1 & S2**). From the forward and backward TMD paths created between the open and closed states, structures are evenly (for every 0.1 Angstrom in the RC) selected to be used for umbrella sampling simulations. In the umbrella sampling simulations from the selected structures along the TMD paths, harmonic restraints are used along the RC. The force constants used in TMD continue are continually used in umbrella sampling simulations (see **SI Table S2**). The biased histograms along the reaction coordinate for each window were unbiased / re-weighted using the weighted histogram analysis method [60]. From the generated biased trajectories a short set of data is removed from the beginning of each for equilibration (10 ns for RTP simulations and 20 ns for ATP simulations) and not used in the construction of the PMF. Overlap for each set of windows was checked along the reaction coordinate (see **Fig S7**). The unbiased probabilities and then the free energy are also calculated using WHAM package [61], following equations:

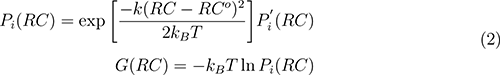

Where *P_i_*(*RC*) and 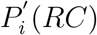 are the unbiased and biased probabilities sampled for the i-th window, respectively. The harmonic restraint potential is shown by 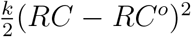 where *RC^o^* is for the initial structure obtained from the TMD insertion path. Finally free energy profile *G* along the RC is calculated taking the logarithm of unbiased probabilities, which represent the PMF.

While constructing the PMF using WHAM, bootstrapping error analysis [62] is used to estimate errors. Bootstrapping re-samples *RC_i_* in each window; from each bootstrapped trajectory *RC_b,i_*(*t*) a new histogram (*h_b,i_*(*RC*)) is created. From the new histograms the PMF and *G_b_*(*RC*) are reconstructed, this process is repeated N times (N = 500 used in this study) generating N bootstrapped PMFs *G_b,j_*(*RC*)(*j* = 1, 2,…, *N*). The uncertainty of a PMF is estimated by a standard deviation calculated by the N bootstrapped PMF’s.

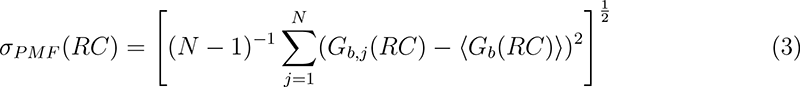

##### Hydrogen Bond Analysis

To examine the corresponding nucleotide insertion dynamics (with intermediate or transition state over-sampled in the umbrella sampling simulations) a hydrogen bond (HB) analysis was performed on the trajectories used to construct a PMF. This was done by taking the last 10 ns of each window and combining them into a single trajectory. The distances for potential HBs were calculated using the MDAnalysis python package. [63] From this analysis, plots were created to indicate when a particular HB was present or not along the windows from open to close along the reaction coordinate. Using a similar formalism salt bridge plots for the NTP polyphosphate were also constructed (see **Fig S8**).

## 4 Results

Upon modeling the active-site closed state complexes for the RDV-TP and ATP insertion, respectively, and then constructing the active-site open state complex to allow the substrate to bind initially (see **SI Methods**), we conducted equilibrium all-atom MD simulations for the closed and open complex systems, bound with RDV-TP or ATP (see **Fig 2**). Base pairing between RDV-TP or ATP with the +1 template nt (Uracil) can well be maintained in the closed state (see **Fig 2** and **Fig S9**). In the open state the base pairing between the RDV-TP or ATP with the template nt appears less or slightly less stabilized. Interestingly, base stacking configuration between RDV-TP and the template nt can be frequently captured (see **Fig S9D&E**), in which the nt base usually stacks upstream relative to RDV-TP (see **Fig 2**). Then we performed TMD simulations to generate the nucleotide substrate (ATP and RDV-TP) insertion paths, and finally conducted a series of umbrella sampling simulations to obtain the nucleotide insertion PMFs for individual systems. The results show uniformly that the closed insertion state is more stabilized than the open initial binding state for each substrate, while the relative stability of the open states (Δ*G^OC^* = *G_open_* – *G_closed_*) and the insertion barriers (Δ*H^ins^* = *G^Barrier^* – *G_Open_*) vary for individual systems. We illustrate results on these systems below, for the ATP insertion, (i) excluding and (ii) including +1 template nt in the RC, initiated from the open state, with ATP base pairing with the template nt; for the RDV-TP insertion, initiated similarly from the (iii) RDV-TP base pairing with the template nt under forcing (i.e., included in the RC), and then from a varied initial configuration, i.e., RDV-TP stacking with the template nt, as the nt (iv) included and (iv) excluded in the RC (i.e., with and without forcing).

**Figure 2:**
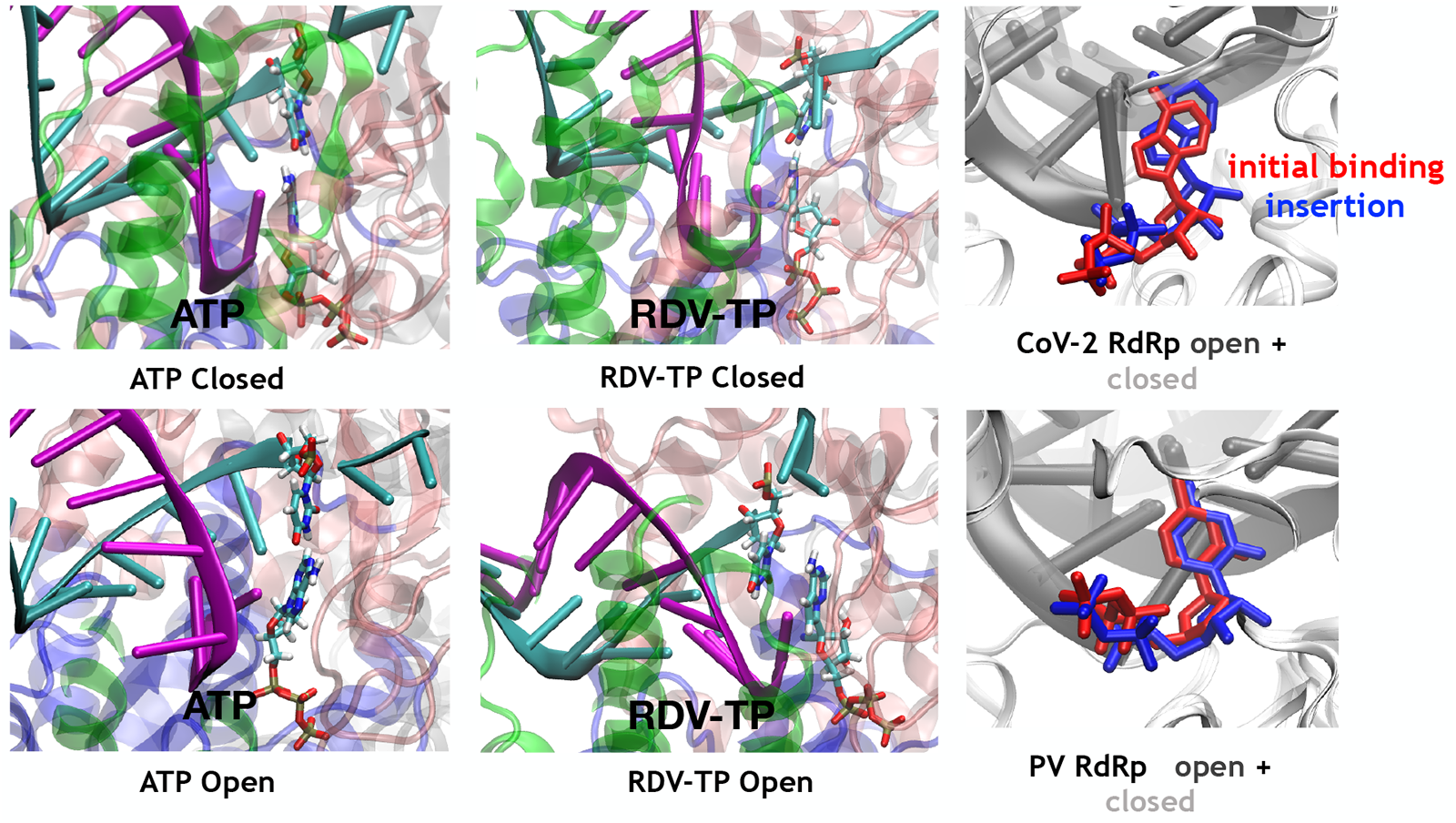
Modeled insertion structural complexes of SARS-CoV-2 for RDV-TP and ATP. Left and Center: The active site views with inserted ATP and RDV-TP shown at the end of equilibrium simulations for the insertion (top) and initial binding (bottom). Right: The open and closed RdRp structures aligned (top), with ATP initial binding and inserted, respectively. The CoV-2 RdRp is shown in comparison with previously studied RdRp from Poliovirus (PV) (bottom) (PDBs: 3ola and 3ol7) [29]

### 4.1 Insertion of ATP into the active site can be facilitated by base pairing with the stabilized template nt (+1)

Upon MD equilibration of the initial binding open-state RdRp complex with ATP (~100 ns; see **SI Fig S10** for RMSD), we found that ATP shows primarily the base pairing initial binding configuration with the +1 template nt. The base pairing interactions seem to be much stabilized in the closed-state ATP insertion configuration (see **Fig 2** left and **SI Fig S9A&B**). By obtaining quasi-equilibrated reference structures from the open-state ATP binding and closed-state ATP insertion complexes, we performed the TMD simulation between these two reference structures and constructed the ATP insertion path for conducting the umbrella sampling simulations. The convergence of the PMFs for the ATP insertion requires about 100~200 ns MD simulation for individual simulation window (see **Fig S11A**). In the first simulation system, ATP constantly forms base pairing with the +1 template nt in the initial binding or active-site open state. We conducted the umbrella sampling simulations by forcing atoms from motif A-G in the palm subdomain and ATP along the TMD insertion path. In this case, the +1 template nt is excluded from the RC, so it is subject only to thermal fluctuations but not the umbrella forcing or constraining. Under such conditions, we noticed that ATP can become highly destabilized (under forcing) by occasionally shifting its base far from the active site in the open state and during barrier crossing (see **Fig 3B&D**). Overall, the ATP insertion can still proceed toward the comparatively stabilized closed state, with ATP base pairing with the template nt much better than in the open state. Overall, the ATP insertion can still proceed toward the comparatively stabilized closed state, with ATP base pairing with the template nt much better than in the open state. Correspondingly, the open to closed free energy drop is obtained as Δ*G^OC^* ~4.8±0.3 kcal/mol and the ATP insertion barriers appears high as Δ*H^ins^* ~5.0±0.3 kcal/mol. During insertion, one can see that motif F-K551 (R555) and K798 (near motif D C-term) constantly form HB interactions with the triphosphate of ATP throughout the process, along with motif F-K545 and the template; motif C-S759 and D760 form HBs with the ATP sugar at open state to barrier crossing, but not into the closed state; S682 (near N-term of motif B) forms transient HB with the template nt during the barrier crossing; motif C-N691 and motif A-D623 form no HBs with ATP sugar until the closed state or crossing the barrier, along with motif F-R553 with the ATP phosphate and motif G-K500 with the template backbone.

**Figure 3:**
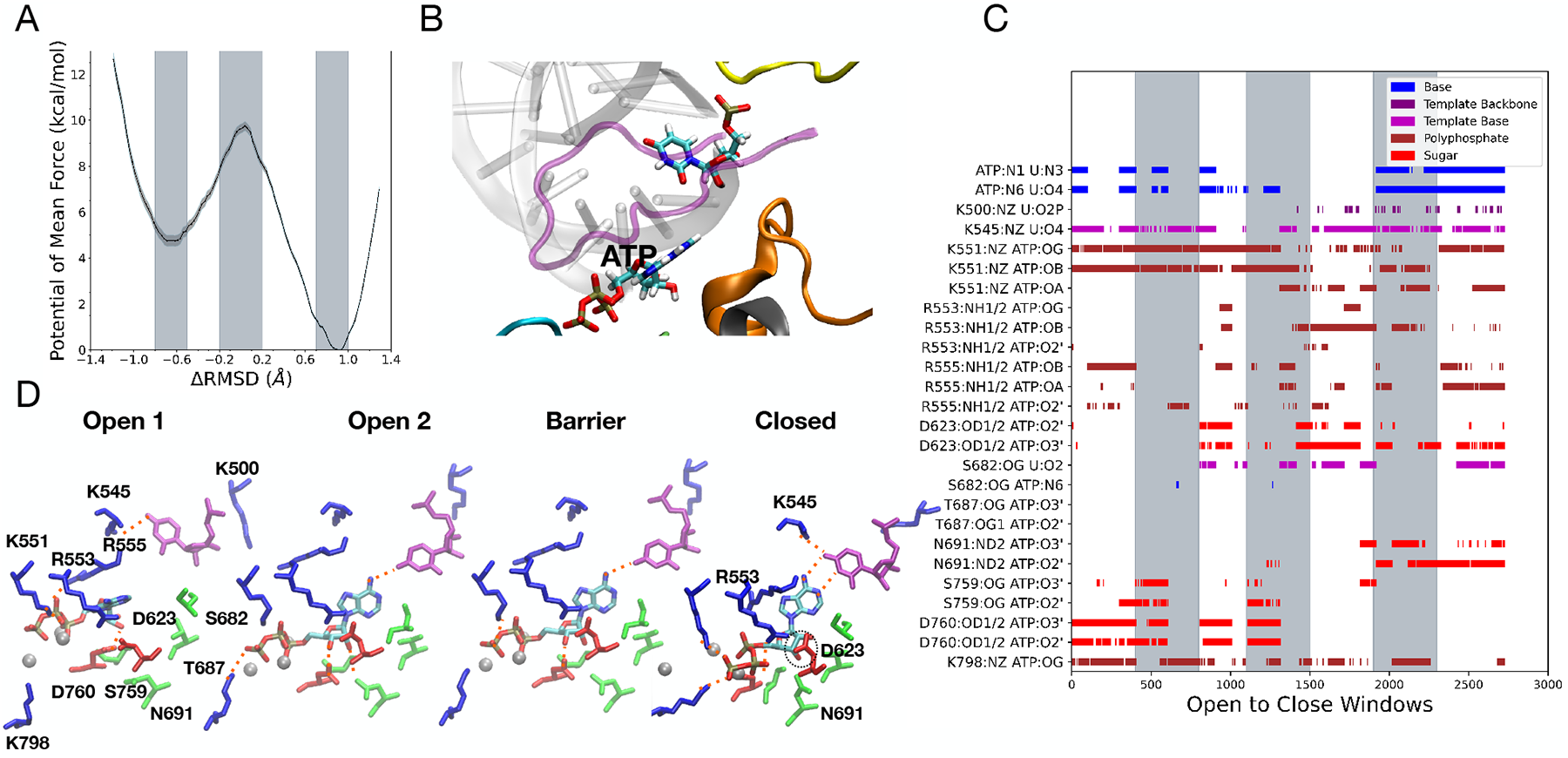
ATP insertion from umbrella sampling MD simulation (without force on the +1 template nt in the RC). **A** PMF with barrier 5.0±0.3 kcal/mol and an initial binding stability of 4.8±0.3 kcal/mol. **B** Open minima conformation with ATP not forming hydrogen bonds with +1 template base. **C** Systematical HB patterns; the grey bars represent Open, Barrier, and Closed regions of the simulation windows (see **SI Fig S8A** for salt bridges). **D** Interaction snapshots from simulation windows: Two open states are shown due to the volatility of the open minima, ATP often flips out of plane from the +1 template base. As the barrier is crossed it begins to form consistent base pairing with the template. Dotted orange lines highlight essential HB interactions.

Next, in order to stabilize the ATP insertion process, we included the +1 template nt in the RC (i.e., with the umbrella forcing) and constructed the second PMF (see **Fig 4**). Consequently, with the ATP:template base pairing better stabilized the ATP base deviated less frequently and not that far from the active site in the open to the barrier crossing state, and ATP base pairing with the template nt can recover sooner after barrier crossing. Notably, the ATP insertion barrier lowers to Δ*H^ins^* ~2.6±0.3 kcal/mol, although the initial open state stability maintains similarly as in the first case (or slightly more destabilized: Δ*G^OC^* ~5.1±0.2 kcal/mol relative to the closed state). Hence, forcing on the template nt or quenching the fluctuations seems to facilitate the ATP insertion, likely by stabilizing the transition state with the ATP-template nt base paring. Such an operation can mimic the spontaneous ATP insertion process that happens sufficiently slowly (e.g. over milliseconds) Overall, the ATP local interactions with nearby amino acids around the active site appear similarly in the two simulation systems, except that in the current template forced condition, the HBs from motif A-D623:sugar and motif G-K500 (&S682): template formed a bit earlier in the open state, and motif F-R555 forms HBs with the ATP phosphates more often throughout the process. Hence, the D623-sugar, R555-phosphate, and the K500/S682-template interactions, along with the template forcing on stabilizing the ATP-template nt base pairing seem to contribute to the lowered ATP insertion barrier.

**Figure 4:**
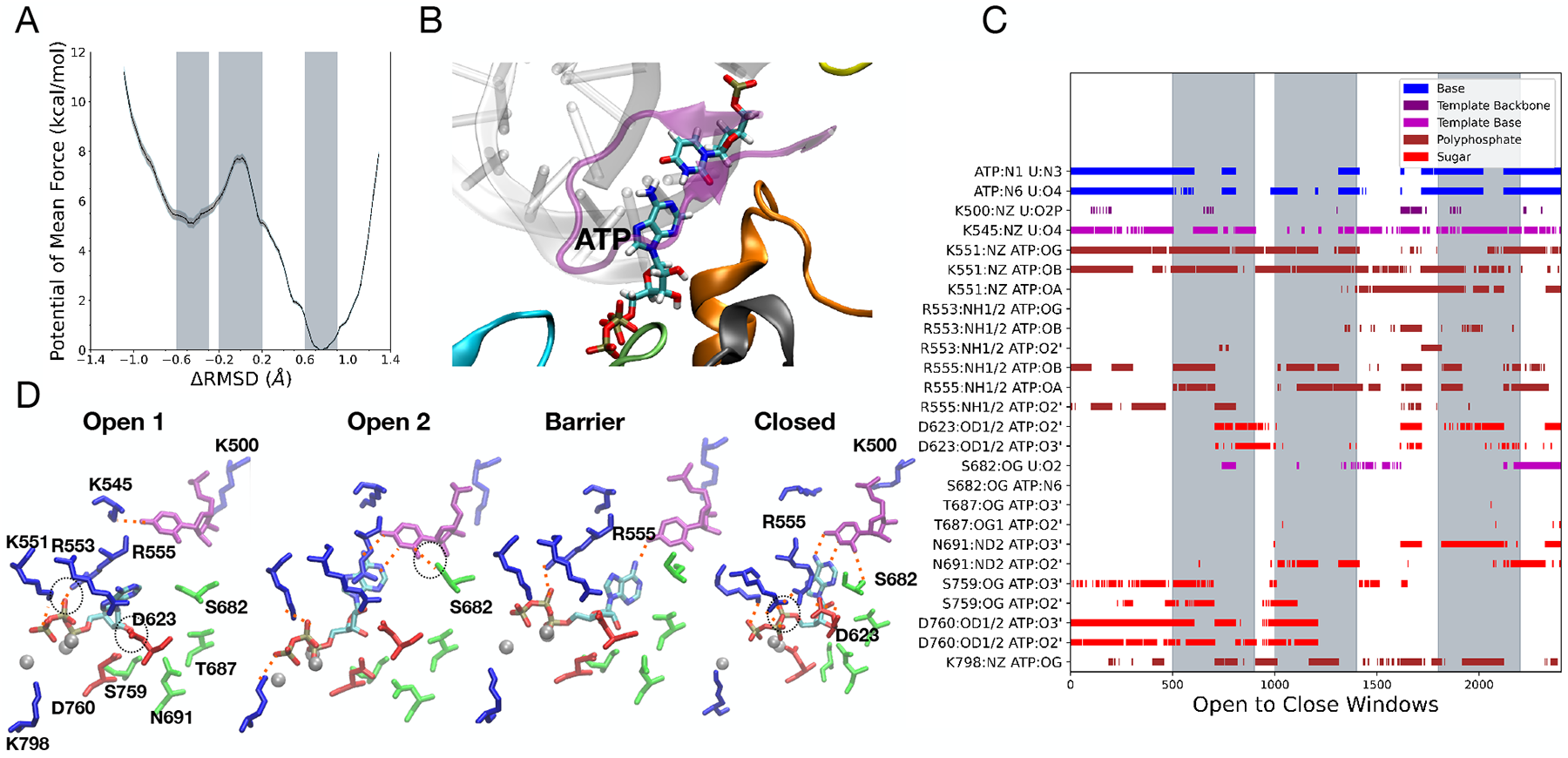
ATP insertion from umbrella sampling MD simulation (with force on the +1 template nt in the RC). **A** PMF with barrier of 2.6±0.3 kcal/mol and an initial binding stability of 5.1±0.2 kcal/mol. **B** Open minima conformation of ATP forming hydrogen bonds with +1 template base. **C** Systematical HB patterns; the grey bars represent Open, Barrier, and Closed regions of the simulation windows as shown in the PMF (see **SI Fig S8B** for salt bridges). **D** Interaction snapshots from simulation windows: Two open states are shown due to the volatility of the open minima. Although ATP still occasionally flips out of plane, it more consistently forms HB with the +1 template base. Dotted orange lines highlight essential HB interactions.

### 4.2 RDV-TP initial stacking with the +1 template nt is more stabilized than the base pairing

Upon MD equilibration of the open-state RdRp complex with RDV-TP (~ 100 ns; see **Fig S12** for RMSD), we found that RDV-TP shows primarily two unique open state binding configurations: one still with standard base pairing and the other with the RDV base stacking with the +1 template base (see **Fig 2** center). We next constructed the PMF for the RDV-TP initially base pairing with the template nt (see **Fig 5**), applying force or constraint to the template (similarly as to the ATP insertion in **Fig 4**). Then we chose the varied initial binding configuration as the RDV forms base stacking with the template nt, keeping the force constraint on the template, and repeated the calculations (see **Fig 6**). Note that the convergences of the RDV-TP insertion energetics happen much faster (~50 ns; see **Fig S11C-E**) than that of the ATP system. The PMF of the RDV-TP insertion starting from the base pairing configuration shows that the insertion barrier is high (Δ*H^ins^* ~5.4±0.3 kcal/mol), comparing to the ATP insertion barrier obtained in the similar conditions (Δ*H^ins^* ~2.6±0.3 kcal/mol from **Fig 4**). The relative stability of the open initial binding state of RDV-TP to the closed insertion state is also measured (Δ*G^OC^* ~4.5±0.3 kcal/mol), slightly lower than that in the corresponding ATP base pairing system (Δ*G^OC^* ~5.1±0.2 kcal/mol from Fig 4). Now motif F-K551, R553&R555 form HBs more or less with the triphosphate of RDV-TP throughout the process, along with motif F-K545 and S682 with the template, and motif C-S759 with the sugar; D760 and motif A-D623 form HBs with the sugar at open state to barrier crossing, not into the closed state; motif C-N691 and motif B-T687 barely forms HB with the sugar until the barrier crossing, along with motif F-K500 and the template. Overall, motif F-R553 and R555 form stronger interaction with the RDV-TP triphosphate than in the ATP insertion cases, while motif A-D623 barely forms HB with the RDV-TP sugar into the closed or insertion state (but with ATP sugar in the insertion state). In contrast, motif B-T687 forms HB with the RDV-TP sugar in the insertion state, while there is no HB interaction of it with ATP at all.

**Figure 5:**
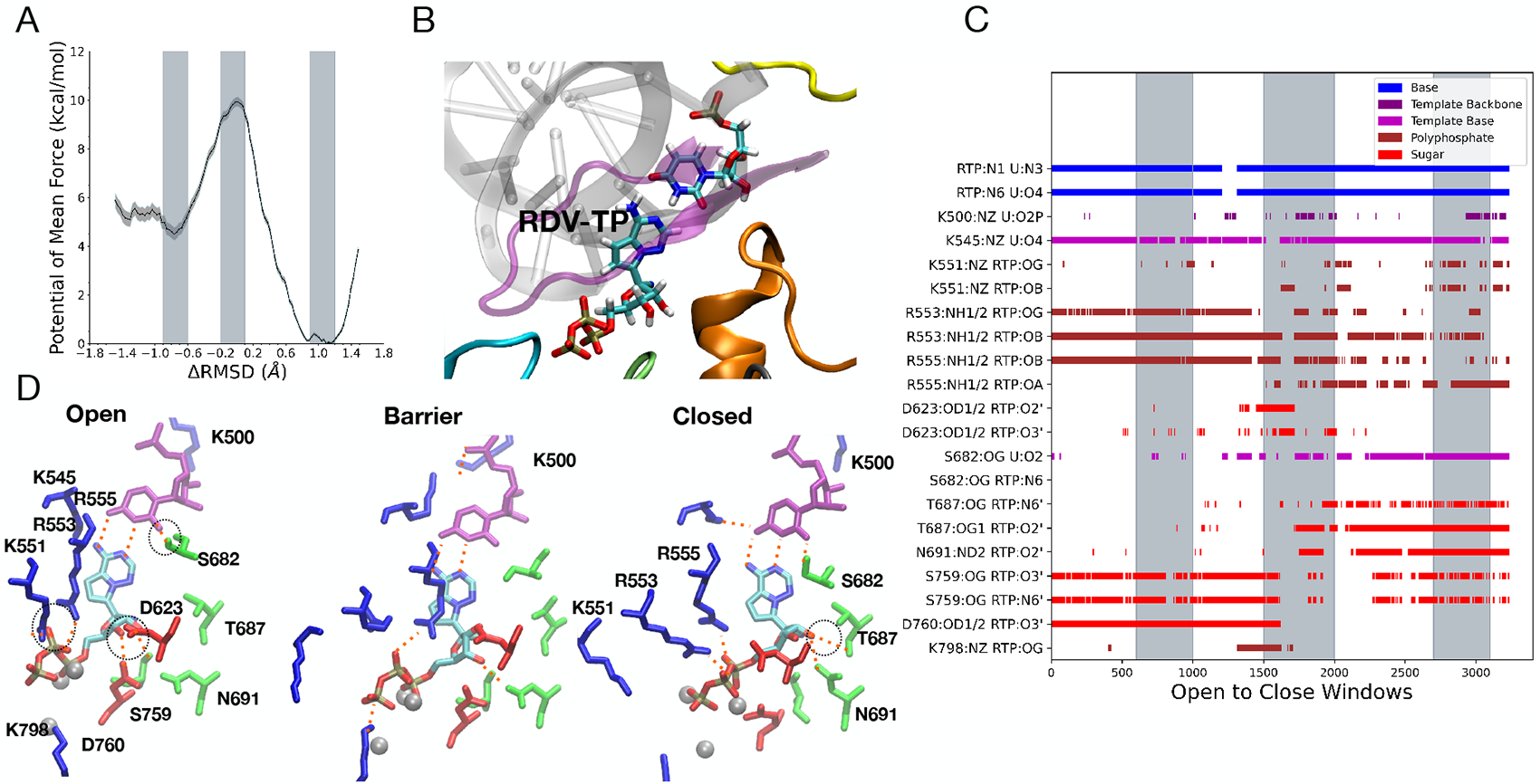
RDV-TP insertion with the open state forming good base pairing from umbrella sampling MD simulation (with force on the +1 template nt in the RC). **A** PMF with barrier of 5.4±0.3 kcal/mol and an initial binding stability of 2.6±0.3 kcal/mol. **B** Open minima conformation of RDV-TP forming hydrogen bonds with +1 template base. **C** Systematical HB patterns; the grey bars represent Open, Barrier, and Closed regions of the simulation windows as shown in the PMF (see **SI Fig S8C** for salt bridges). **D** Interaction snapshots from simulation windows: Throughout the open state stable HB form with the RDV and+1 template base. Dotted orange lines highlight essential HB interactions.

More interesting results come from comparing RDV-TP insertion energetics and interactions simulated at the varied conditions. In **Fig 6**, we show the PMF from RDV-TP initially stacking with the +1 template nt, with forcing still implemented. Although the insertion barrier (Δ*H^ins^* ~5.2±0.3 kcal/mol) remains similarly high as the above case (from **Fig 5**), the relative stability of the initial open state to the final insertion or closed state changes (to Δ*G^OC^* ~2.6±0.2 kcal/mol), indicating that the initial stacking configuration of RDV-TP is more stabilized (about −3~ −2 k_*B*_T) than the initial base configuration with the template nt). By comparing the HB patterns (**Fig 5** and **Fig 6C**), one finds that the stabilizing interactions to the base stacking configuration at the open state mainly come from motif B-N691 and motif A-D623 with the sugar, S682 interaction with the RDV-TP base, K798 (near motif D) along with K551 interaction with the phosphate, as well as motif G-K500 interaction with the template nt. The R555/R553 interaction with the RDV-TP triphosphate weaken somehow from the initial base pairing to the stacking configuration, which appear to matter for the insertion barrier, as demonstrated next.

**Figure 6:**
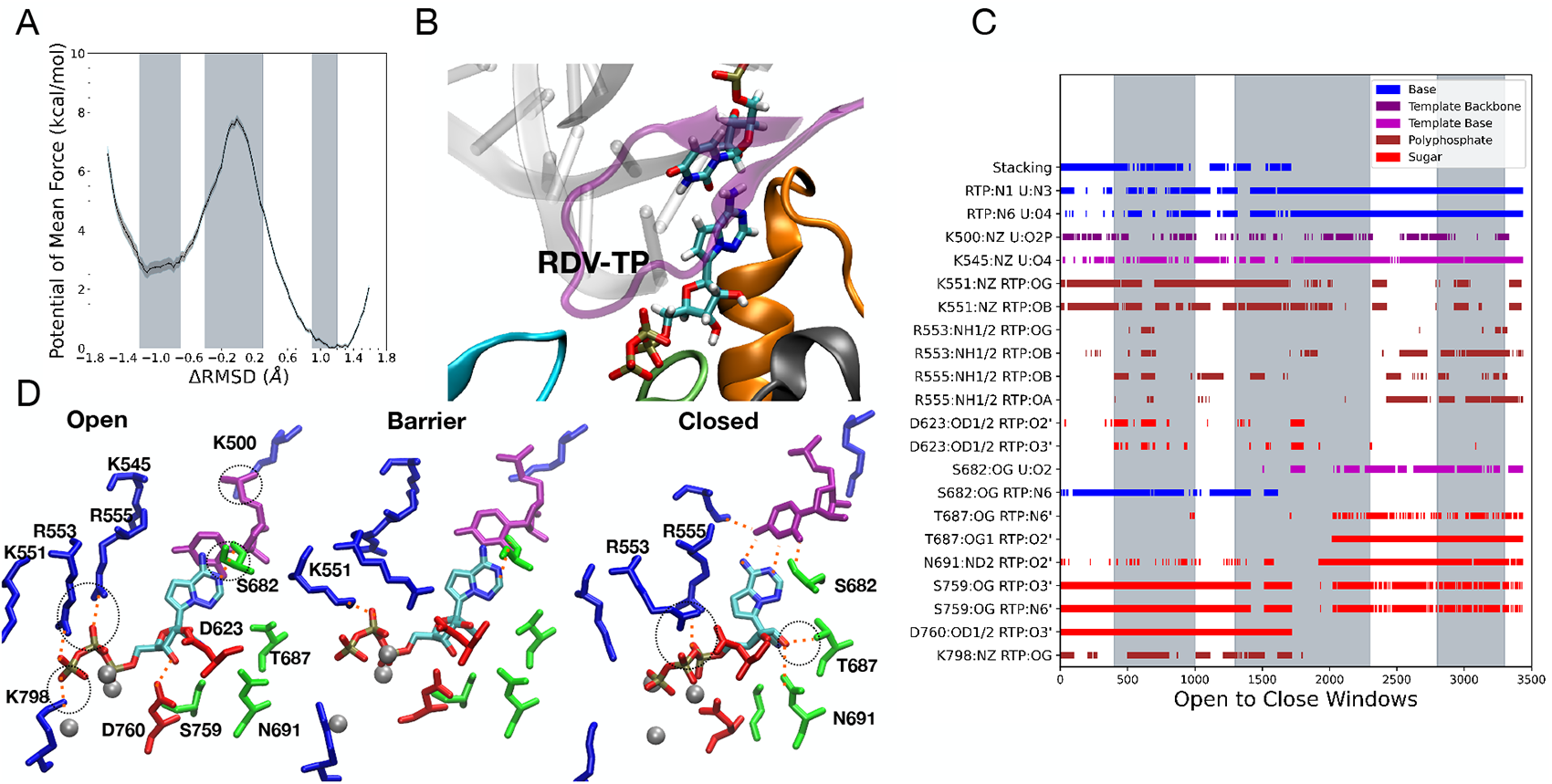
RDV-TP insertion with the open state forming base stacking with the +1 template base from umbrella sampling MD simulation (with force on the +1 template nt in the RC). **A** PMF with barrier of 5.2±0.3 kcal/mol and an initial binding stability of 2.6±0.2 kcal/mol. **B** Open minima conformation of RDV-TP forming base stacking with +1 template base. **C** Systematical HB patterns; the grey bars represent Open, Barrier, and Closed regions of the simulation windows as shown in the PMF (see **SI Fig S8D** for salt bridges). **D** Interaction snapshots from simulation windows: Throughout the open state base stacking forms resulting in a more stable minima. Dotted orange lines highlight essential HB interactions.

### 4.3 RDV-TP insertion to the active site is highly facilitated by non-stabilized phosphate positioning

Since the above results show that the RDV-TP initial stacking with the +1 template nt is more stabilized than the base pairing configuration, we further explored the RDV-TP insertion barrier by removing the forcing on the +1 template nt (i.e. being excluded from the RC). Notably, the insertion now is greatly facilitated by allowing sufficient fluctuations on the template, such that the insertion barrier becomes lowest (Δ*H^ins^* ~1.5±0.2 kcal/mol; see **Fig 7A**). Meanwhile, the relative stability of the open binding state to the closed insertion state of RDV-TP maintains (Δ*G^OC^* ~2.7±0.1 kcal/mol), as in the above system from **Fig 6**).

**Figure 7:**
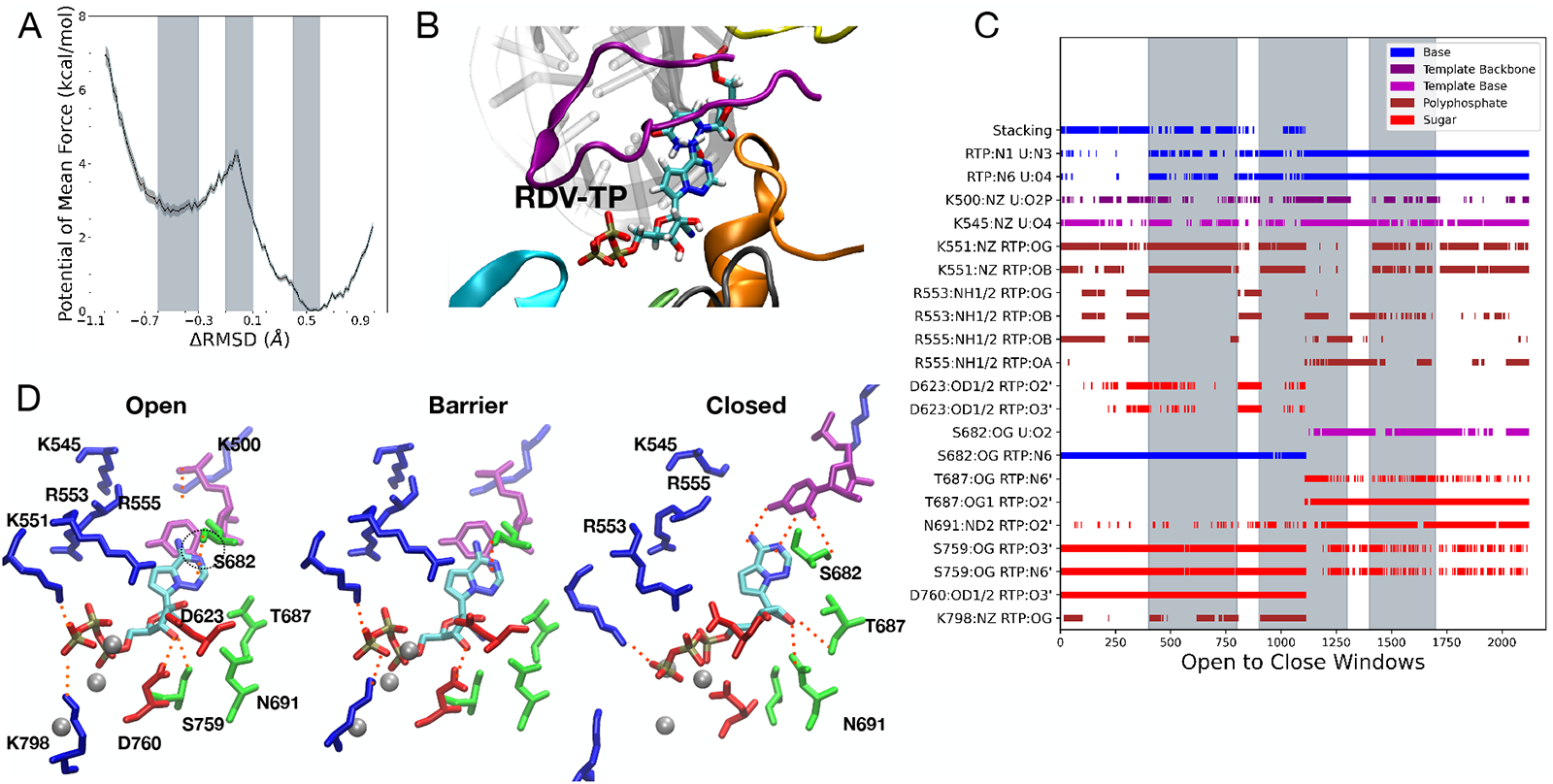
RDV-TP insertion with the open state forming base stacking with the +1 template base from umbrella sampling MD simulation (with no force on the +1 template nt in the RC). **A** PMF with barrier of 1.5±0.2 kcal/mol and an initial binding stability of 2.7±0.1 kcal/mol. **B** Open minima conformation of RDV-TP forming base stacking with +1 template base. **C** Systematical HB patterns; the grey bars represent Open, Barrier, and Closed regions of the simulation windows as shown in the PMF (see **SI Fig S8E** for salt bridges). **D** Interaction snapshots from simulation windows: Throughout the open state base stacking forms resulting in a more stable minima. Dotted orange lines highlight essential HB interactions.

Close examinations show that the RDV-TP stacking association with the template nt can be more stabilized when there is no forcing or constraining on the template nt, while forcing the template nt can indeed cause large deviations between the RDV and template bases. Hence, it appears that thermal fluctuations on the template nt can actually support the RDV base stacking with the template nt along with “shaking” the motif F-R553/R555 interaction off triphosphate before transition toward the insertion configuration, in which RDV-TP can form very stabilized base pairing interactions with the template nt.

Additional close inspections on the RDV-TP local interactions with nearby residues show that the majority of HB and SB interactions are similar between the cases without and with forcing on the +1 template nt (**Fig 7C** and **Fig 6C**). Interestingly, one can identify that both HB and SB interactions from R555 and R553 (located on the motif F) with the triphosphate of RDV-TP, which are formed for the RDV-TP initial binding in the former stacking case with template forcing (see **Fig 6C**), but become absent in the current case (without forcing on the template nt, **Fig 7C**). Otherwise, the local HB/SB interactions with RDV-TP are highly similar for the two systems (**Fig 6** and **Fig 7**), both initiated from the RDV-TP base stacking with the template nt binding configuration. Hence, in the RDV-TP insertion, the presence of the template forcing (or reduced fluctuations) along with the R555 (and R553) interaction with the triphosphate seems to hinder the RDV-TP insertion, which appears to be opposite to the trend in the ATP insertion (i.e., stronger R555/R553-ATP phosphate interaction in the open state under template forcing condition leads to a lowered ATP insertion barrier).

## 5 Discussion

In this work we modeled and simulated insertion of the triphosphate form nucleotide analogue drug remdesivir (RDV-TP) into the SARS-CoV-2 RdRp active site, in comparison with natural nucleotide substrate ATP. Our work is based on high-resolution cryo-EM structures solved for the SARS-CoV-2 nsp12 in complex with cofactors nsp7 and nsp8 [23, 24], modeled in an active-site open form (PDB: 7BTF) for the nucleotide initial binding, and in an active-site closed form (PDB: 7BV2) for the stabilized nucleotide insertion, prior to catalytic addition of the nucleotide to the synthesizing RNA chain. The viral RdRp or nsp12 in the coronavirus species works with other non-structural proteins (nsp7 to nsp16) for viral genome replication and transcription [64, 65], with nsp7 and nsp8 the cofactors to assist the replication machinery stability and processivity along the viral genome, and with nsp13 [66, 67] and nsp14 [68] functioning as helicase and exonuclease, respectively. In the simulation of the nsp12-nsp7-nsp8 complex along with RNA strands, we found that shortening of the nsp8 N-terminal (e.g. to start from residue M67) is necessary to stabilize the simulation complex in all-atom explicit water condition. It is however noted that the two copies of nsp8 can extend very long as ‘sliding poles’ on a protruding exiting RNA duplex, as being captured from another high-resolution cryo-EM complex of nsp12-nsp7-nsp8 [25]. In modeling of an initial binding complex of the nucleotide or analogue, we placed ATP or RDV-TP to the open active site of CoV-2 RdRp, according to RdRp structural alignments between the product complex (closed form) of RDV-TP and the open one. Accordingly, the positioning of RDV-TP or ATP appear similar between the open and closed structures. Molecular docking and simulation equilibration confirmed such an initial nucleotide binding configuration is dominant (see **SI Fig S3&S4**), which also appears similar to that being captured in the polio virus (PV) RdRp [29]. Hence, for the RDV-TP and ATP insertion probed in this work, we focus mainly on subtle local interactions around the active site of the viral RdRp as for the incoming nucleotide being recruited, interrogated, and re-positioned to allow chemical addition. Meanwhile, we note that the open and closed forms of the viral RdRp structure still involve collective movements of the highly conserved motifs (A to G) which we manipulate as a whole in the umbrella sampling simulations, to ensure the concerted nucleotide insertion. Note that motifs A to E are located in the palm subdomain hosting the active site, with motif C mainly responsible for catalysis, and motif A,B, and D for nucleotide binding and selection; motif F-G from in the fingers subdomain also impacts on the incoming nucleotide entry as well as the +1 template nt for the Watson-Crick (or WC) base pairing or fidelity check [15,69].

Correspondingly, we conducted first the equilibrium MD simulations, which show that upon the initial binding, ATP frequently forms the WC base pairing with the template nt but with notable fluctuations; in contrast, RDV-TP primarily forms base stacking with the template nt, squeezing the template base to upstream most of time. Although RDV-TP has also been sampled in base paring with the template nt, such a binding configuration appears stable. In the closed RdRp or insertion state, RDV-TP anyhow forms highly stabilized base pairing with the template nt, with even lower fluctuations than ATP for natural base pairing. APBS mapping zoomed into the closed active site of CoV-2 RdRp shows notable differences between the local electrostatic environment around the inserted RDV-TP and ATP, in particular around the sugar region, where an extra cyano group is attached to RDV-TP, with T687 and N691 associated nearby. In order to see how exactly RDV-TP and ATP insert into the active site from the initial binding state, as the open active site closes, we then performed the TMD and umbrella sampling simulations connecting the open and closed RdRp complex structures, with slightly varied initial and collective coordinate forcing conditions.

The purpose of running the TMD simulations was to construct feasible dynamical paths of the nucleotide insertion to be used in the umbrella sampling simulations for the PMF construction, upon that the structural dynamics (with enhanced sampling in the transition state or barrier region) and energetics (or free energy profiles) of the insertion processes reveal and can be further compared. Our simulations first confirm that the nucleotide inserted or the closed form of the CoV-2 RdRp is indeed much more stabilized than the open form for nucleotide initial binding (about −3 to −5 kcal/mol), for RDV-TP or ATP. While the base pairing configurations of the initial binding ATP and RDV-TP are similarly stabilized (~5 kcal/mol) relative to the corresponding closed insertion state, such an initial binding configuration is only dominant to ATP but not RDV-TP. Essentially, our calculations show that RDV-TP primarily forms base stacking with the +1 template nt rather than base pairing upon initial binding. Comparison between the RDV-TP insertion simulations conducted with varied initial binding configurations (stacking and base pairing) shows that motif A-D623 and motif B-N691 particularly stabilize the RDV-TP base stacking over the base pairing in the open state, by forming HBs with the sugar; motif B-S682 forms HB contact with the RDV-TP base, only in the base stacking configuration; motif F-K551 and K798 near the C-terminal of motif D stabilize the base stacking configuration by forming HB (or SB) interactions with the RDV-TP triphosphate, along with motif G-K500 interacting with the + 1 template backbone. The stabilization leads to ~-2 kcal/mol (or ~-3 kBT) relative initial binding free energy between the RDV-TP stacking and base pairing configuration. A docking stabilization energetics (~-0.6 kcal/mol) between the RDV-TP and ATP was reported to a homology modeled CoV-2 RdRp [19], and a similar energetic score revealed from our own docking trials (using the open form RdRp complex with RNA strands, see **SI Fig S3&S4**). Hence, it seems that the initial binding of RDV-TP to the CoV-2 RdRp can be about −2 to −3 kcal/mol more stabilized than ATP. An alchemical MD simulation for relative binding free energy calculation have presented a comparable stabilization energetics between RDV-TP and ATP (~-2.8 kcal/mol) upon binding to the RdRp active site [20]. Nevertheless, the alchemical calculation was conducted in the absence of RNA, so it is unable to be compared in regard to the template RNA configuration. The computational results so far consistently point out that RDV-TP can bind to the CoV-2 RdRp active site (in an open form) more favorably than the natural nucleotide substrate ATP.

Nevertheless, the initial binding to RdRp only provides an initial nucleotide association and selection checkpoint to the nucleotide addition cycle or NAC. The followed insertion of the nucleotide to the active site becomes a next and likely the most important checkpoint in the NAC, in particular, for the single-subunit handlike RNA or DNA polymerases (RNAPs or DNAPs). In several such polymerase species, the nucleotide insertion is rate-limiting (or partially rate-limiting) [33,70,71], thus being critical for nucleotide selection [34]. Comparing to phage T7 RNAP we studied previously [35, 36], for which a substantial fingers subdomain rotation happens with respect to the palm subdomain (from open to closed) during the nucleotide insertion, the viral RdRp conformational changes in accompany with the nucleotide insertion are mainly the active site distortions (from open to closed) [29], though remote residues on the structural motifs (A-G) can be more or less involved in the process. From the TMD simulations enforcing the CoV-2 RdRp from open to closed (see **SI Fig S6**), we found that the motif A and D close similarly as that in PV RdRp [30]. Interestingly, the inserting ATP or RDV-TP has the base easily re-positioned toward the closed configuration in the TMD simulation, but has the triphosphate moiety hardly reaching to the targeted closed configuration (see **SI movies S1 & S2**). Hence, re-positioning of the triphosphate during the nucleotide insertion appears to link to events of crossing the free energy barrier. In current umbrella sampling simulations, the ATP or RDV-TP insertion barrier indeed depends on the relative template nt configuration or fluctuations, as well as local residue interactions with the triphosphate. In the ATP insertion, a comparatively low energetic barrier (~2.6 kcal/mol) shows when the template nt is enforced or constrained to maintain stabilized base paring with ATP as if in the long-time unperturbed nucleotide insertion; the motif F-R555 interaction with the ATP phosphates along with motif A-D623 interaction with the sugar at the open state seems to facilitate the further ATP insertion. In comparison, for RDV-TP, the insertion barrier can be even lower (~1.5 kcal/mol) when it is inserted without enforcing the template nt, so that the initial base stacking between RDV-TP and template nt can proceed freely to easily transit to the base pairing configuration into the closed insertion state. Contrary to the ATP insertion, motif F R555/R553 close interactions (hydrogen bonding and salt-bridge) with the RDV-TP triphosphate in the open state appears to impede the RDV-TP insertion, which happens as the template nt is enforced in the simulation, no matter which initial configuration RDV-TP starts with the template (base stacking or pairing). Current simulations comparing RDV-TP and ATP thus suggest that the nucleotide insertion is coordinated by +1 template nt as well as some interactions on the nucleotide upon initial binding, in which the triphosphate stabilization and re-positioning appear to be essential. It should be pointed out that the triphosphate reorientation of the incoming nucleotide had been suggested for the PV RdRp fidelity control [72]. Additionally, it is interesting to notice that motif F-R555 structurally corresponds to R174 from PV RdRp and R158 in HCV RdRp [15], as well as to Y639 from T7 RNAP that is key to nucleotide selectivity and polymerase translocation [35]. Overall, the ATP insertion seems to be facilitated by an insertion path with quenched fluctuations on the +1 template nt for stabilized base pairing, while the RDV-TP insertion dominated by the template base-stacking populations, is supported by freely fluctuating template nt, leading to transition to the highly stabilized base pairing configurations, with an insertion free energy barrier as low as ~1.5 kcal/mol or ~2-3k_*B*_T, marginally above thermal fluctuations.

Both the inserted ATP and RDV-TP can be then further stabilized well in the active site by the base paring interaction with the template nt. Though we haven’t yet conducted energetic calculations to evaluate the relative stability between the RDV-TP and ATP in the insertion state, the equilibrium simulations of the insertion complexes of the two species suggest that the RDV-TP can be similarly or even more stabilized than ATP in the closed insertion state. There are also specific interactions that can well distinguish the natural nucleotide substrate from the nucleotide analogue in the insertion state: motif A-D623 forms specific HB contact with the ATP sugar but not with the inserted RDV-TP; K798 near motif D also closely with the ATP gamma-phosphate into the insertion state, but not closely with that of RDV-TP; in contrast, motif B-T687 specifically forms HB with the RDV-TP sugar but not with that of ATP. The overall results (see **Table 1**) thus suggest that binding/insertion of RDV-TP can be more facilitated than the natural substrate ATP to the active site of SARS-CoV-2 RdRp, seemingly consistent with in vitro measurements of the Michaelis-Menten constant K_*m*_ ~ 0.0089 *μ*M and 0.03 *μ*M obtained for RDV-TP and ATP, respectively [9]. If the nucleotide insertion is a single rate-limiting step (i.e., as in T7 RNAP), then V_*max*_ should also be significantly larger for RDV-TP than that for ATP, due to a lowest insertion barrier of the RDV-TP. However, the *in vitro* measurements of V_*max*_ are similar for RDV-TP and ATP [9]. Hence, other rate-limiting steps than the pre-chemical NTP insertion can exist in the NAC of the CoV-2 RdRp, e.g., the chemical catalysis, which may happen a bit slower for RDV-TP than ATP, so that overall the maximum elongation rates become similar for the two nucleotide species. More close examinations of stepwise kinetics of SARS-CoV-2 RdRp are therefore expected, ideally for both cognate and non-cognate nucleotide species, so that substantial information on the complete NAC as well as nucleotide selectivity could reveal. Note that following a successful RDV-TP incorporation to the end of viral RNA chain, additional nucleotide insertion still appears viable until the addition of the nucleotide downstream +3 to the incorporated RDV analog. Such mechanism has been suggested as a delayed chain termination of the nucleotide analogue [9], which arises likely due to aberrant impacts of incorporated analogue on the synthesizing RNA chain in association with the viral RdRp, together with failure of ExoN cleavage or proofreading to the nucleotide analogue.

**Table 1.**
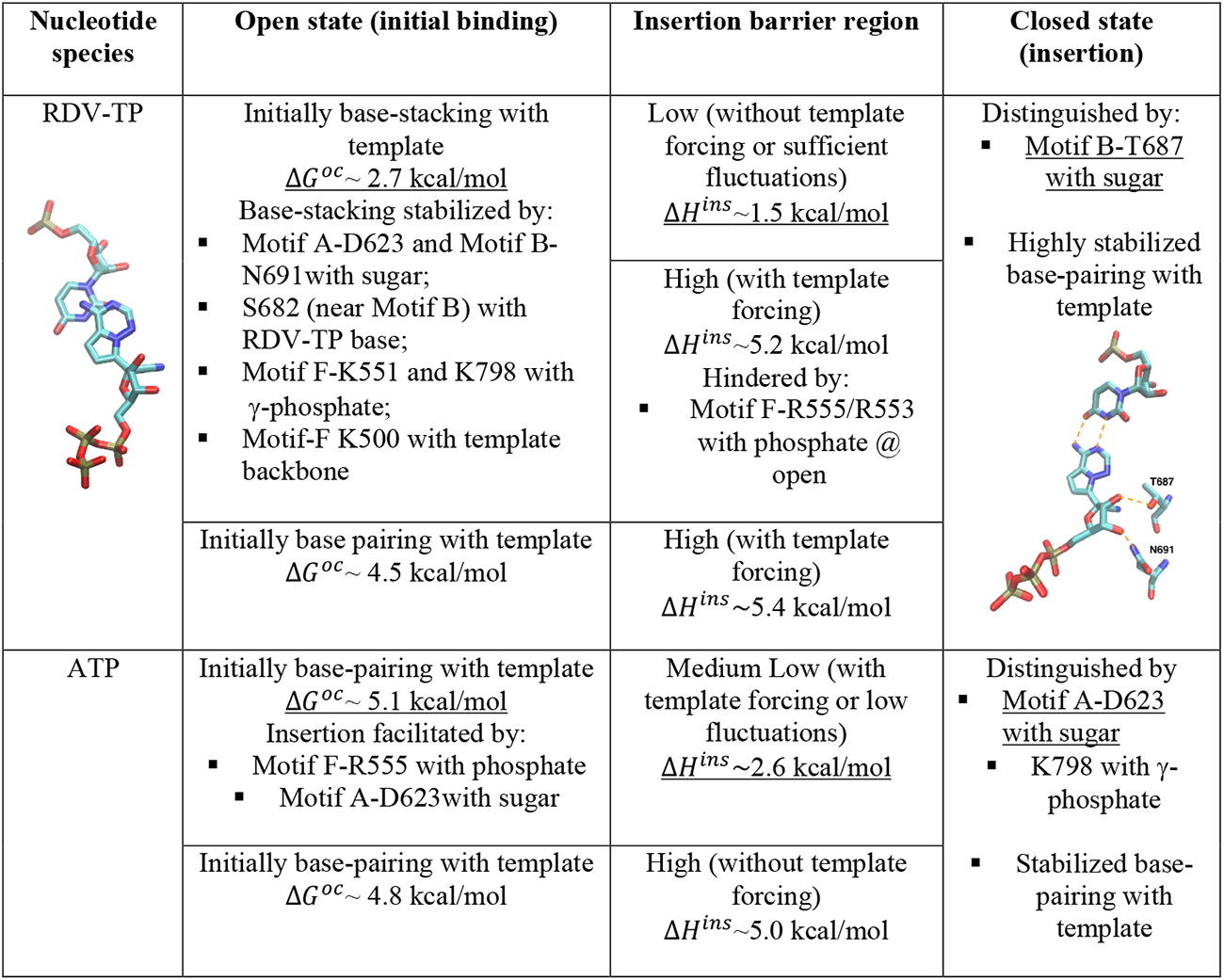
Summary of essential interactions and energetics during RDV-TP and ATP insertion

## 6 Conclusions

Via modeling and all-atom MD simulation, we found that remdesivir nucleotide analogue can bind to the open active site of SARS-CoV-2 RdRp via base stacking with the +1 template nt. Such a stacking configuration appears to be more stabilized than the Watson-Crick base pairing configuration formed between ATP and the template. Umbrella sampling simulations further show that the remdesivir analogue stacking with the fluctuating template then inserts or transits to form high-stabilized base pairing with the template as the active site closes. The corresponding insertion barrier for remdesivir analogue is even lower than that of a low-energetic path of the ATP insertion, during which the template forms stabilized in base pairing with ATP. Additionally, our analyses on hydrogen bonding and salt-bridge interactions during the nucleotide or analogue insertion show that (i) the initial remdesivir base stacking can be particularly stabilized by motif B-N691 with sugar, S682 with base, and motif F-K500 with the template; (ii) insertion of remdesivir analogue can be facilitated by thermal fluctuations but hindered by motif-F R555/R553 interaction with the triphosphate, while insertion of ATP is made easier by lowering fluctuations and taking advantage of the R555/R553 interaction with the triphosphate; (iii) the inserted remdesivir analogue and ATP are distinguished by specific sugar interaction via motif B-T687 and motif-A D623, respectively. Such findings also reveal potential SARS-CoV-2 RdRp fidelity control via particular residue interactions with the nucleotide substrate sugar, base, and triphosphate moieties, along with +1 template coordination.

## Conflicts of interest

There are no conflicts to declare.

## Acknowledgements

Current work is supported by National Science Foundation (NSF) Award #2028935. This work used resources services, and support provided via the COVID-19 HPC Consortium (https://covid19-hpc-consortium.org/), which is a unique private-public effort to bring together government, industry, and academic leaders who are volunteering free compute time and resources in support of COVID-19 research. This research used resources of the Oak Ridge Leadership Computing Facility, which is a DOE Office of Science User Facility supported under Contract DE-AC05-00OR22725. JY has also been supported by the CMCF of UCI via NSF DMS 1763272 and the Simons Foundation grant #594598 and start-up funding from UCI. CL is supported by National Natural Science Foundation of China grant #12005029 and Natural Science Foundation of Chongqing Grant #cstc2020jcyj-msxmX0811 and the Start-up Founding of Chongqing University of Posts and Telecommunication (A2020-029).

## Notes

### Competing Interest Statement

The authors have declared no competing interest.

### Summary of Updates

All figure captions revised due to formatting error.

