## Supplementary material for "Probing remdesivir nucleotide analogue insertion to SARS-CoV-2 RNA dependent RNA polymerase in viral replication": SARS_Cov_2_Paper_SI_5_compressed.pdf

### Supplementary Information

#### Methods

##### Modeling the substrate insertion complex of RdRp

Since a tertiary elongation complex of RdRp (with a full length nsp12) was captured together with RNA strands (template and primer) and the RDV analog incorporated (post-catalysis or product state; PDB: 7BV2), presumably in the active-site closed state, we built an RDV-TP insertion model (pre-catalytic) of the CoV-2 RdRp directly using this tertiary complex, only replacing the incorporated RDV analog at the 3'-end of the RNA primer strand by a pre-catalytic RDV-TP.

###### Missing Residues Added

PDBID:7btf is missing the following residues in the N-terminus domain: 1-30, 51-68, 75, 103-111, 895-906, and the following in the thumb sub-domain 920-932. Missing residues were completed using MODELLER 9.24 [1] with the apo structure PDBID:6M71 which is only missing residues 1-4.

###### Protonation (details)

Propka and pdb2pqr used to predict protonation states of Histidine residues:

- HID: 75 99 113 133 256 347 355 362 439 572 599 613 810 816 882 898
- HIE: 82 295 309 381 642 650 725 752 872 892

- Residue 295 and 642 are manually selected due to their orientation with Zn ion such that the proton is not oriented near the metal.

#### Constructing the initial biding complex and docking

The active-site open structure of the CoV-2 RdRp for NTP initial binding was obtained from the first determined cryo-EM structure (PDB: 7BTF). Then RDV-TP was placed to the active site of the open state structure (along with the RNA template and primer strand) by aligning the RdRp structure from the tertiary RDV-TP insertion complex (the closed one constructed above) with that of the open one, and then shifting the RDV-TP and RNA strands from the tertiary complex to the open state structure accordingly. Followed, the modeled structural complex would be subject to MD simulation equilibration.

In order to test whether the above constructed initial binding complex was reasonable, we also performed docking of RDV-TP (or RTP below) and ATP as ligands onto the open structural complex of RdRp (nsp12+nsp7+ns8 and RNA together as the receptor), using AutoDock Vina software [2]. The receptor complex was prepared by deletion of water molecules, addition of hydrogen molecules and by computing Kollman charges. The ligands (RTP and ATP) were prepared by computing Gasteiger charges. A grid box (x=40 Å, y=40 Å, z=40 Å) is specified around the active site for the search space on the receptor within which various positions of the ligand are to be considered. An energy range of 4 and exhaustiveness of 8 were assigned. Conformations with lowest binding free energetic scores are considered most stable or optimal.

#### RDV-TP Force Field

##### Atom Types

In general the majority of atom types were kept the same as the adenosine where possible from the amber force field. [3]. The swapped and additional atoms (nitrile functional group) used the following atom types for RDV: C9:CK, C7:CK, N4:na, C6':c1, and N6':n1. Were the lower case atom types are taken from the Generalized Amber Force Field (GAFF) [4].

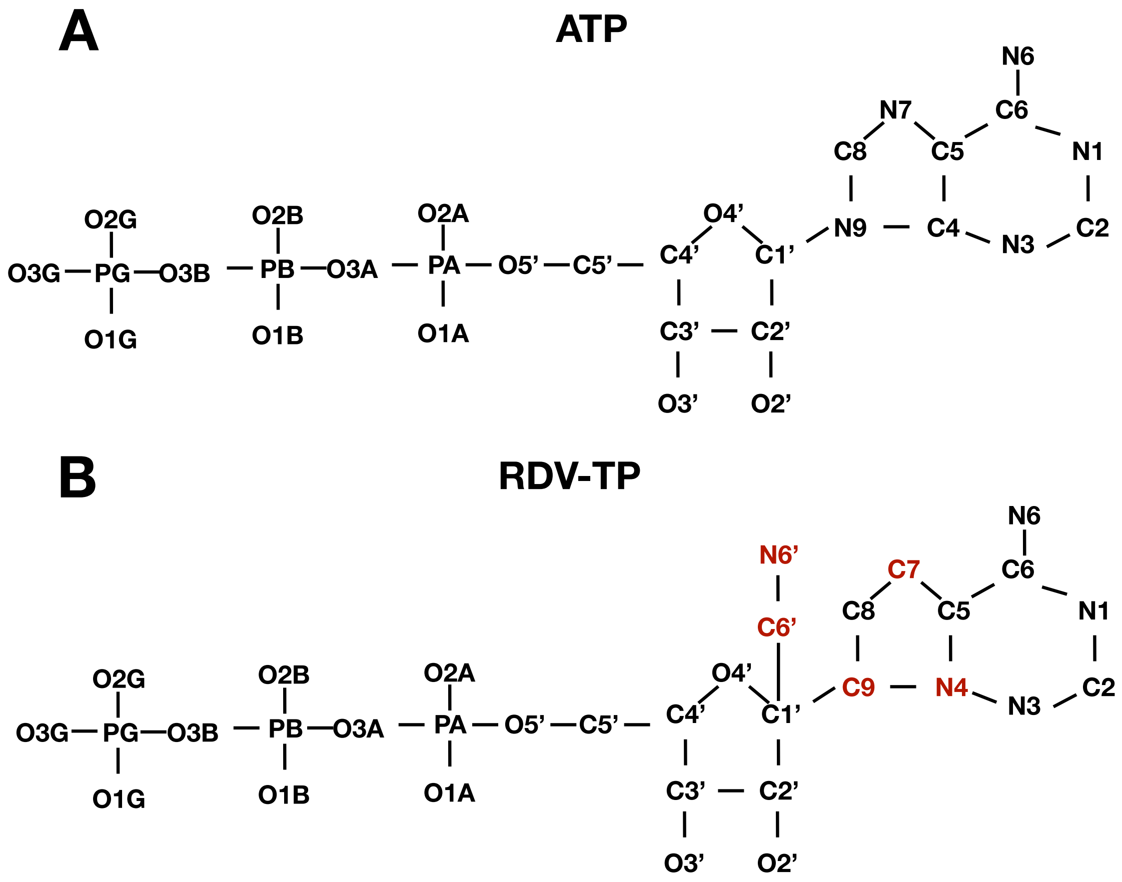


Figure S1: Comparing **A.** Adenosine Triphosphate and **B.** Remdesivir Triphosphate heavy atom molecular structures. Hydrogens are omitted for a clear representation of the heavy atoms. Atoms colored in red highlight the key differences in RDV which require selection of different atom types. For partial charge calculation the RDV-TP is truncated at the O5' with the addition of a hydrogen H5T. H3T is the hydrogen atom bonded to the O3' oxygen.

##### Partial Charges

During Restrained Electrostatic Potential method (RESP [5]) the 3' 5' hydroxyl atomic charges are constrained [6] to O5'= -0.6223e, H5T=0.4295e, O3'=-0.6541e, H3T=0.4376e, partial atomic charges for the truncated Remdesivir are generated.

Table S1: Summary of Partial Charges used for the RDV-TP force field compared with ATP. Charges are separated by section of the NTP.

| Atom (PolyP) | ATP | RTP | Atom (Sugar) | ATP | RTP | Atom (Base) | ATP | RTP |
|---|---|---|---|---|---|---|---|---|
| O5’ | -0.59870 | -0.59870 | C5’ | 0.05580 | 0.039981 | N9 | -0.02510 | N.A. |
| PA | 1.25320 | 1.25320 | H5’1 | 0.06790 | 0.085276 | C9 | N.A. | -0.118619 |
| O1A | -0.87990 | -0.87990 | H5’2 | 0.06790 | 0.085276 | C8 | 0.20060 | -0.228326 |
| O2A | -0.87990 | -0.87990 | C4’ | 0.10650 | 0.083427 | H8 | 0.15530 | 0.199975 |
| O3A | -0.56890 | -0.56890 | H4’ | 0.11740 | 0.065203 | N7 | -0.60730 | N.A. |
| PB | 1.38520 | 1.38520 | O4’ | -0.35480 | -0.332867 | C7 | N.A. | -0.259805 |
| O1B | -0.88940 | -0.88940 | C1’ | 0.03940 | 0.130365 | C5 | 0.05150 | -0.394730 |
| O2B | -0.88940 | -0.88940 | H1’ | 0.20070 | N.A. | C6 | 0.70090 | 1.014028 |
| O3B | -0.53220 | -0.53220 | C6’ | N.A. | 0.461023 | N6 | -0.90190 | -1.042464 |
| PG | 1.26500 | 1.26500 | N6’ | N.A. | -0.505959 | H61 | 0.41150 | 0.443695 |
| O1G | -0.95260 | -0.95260 | C3’ | 0.20220 | 0.329872 | H62 | 0.41150 | 0.443695 |
| O2G | -0.95260 | -0.95260 | H3’ | 0.06150 | 0.076195 | N1 | -0.76150 | -0.863108 |
| O3G | -0.95260 | -0.95260 | C2’ | 0.06700 | -0.074121 | C2 | 0.58750 | 0.630021 |
| | | | H2’1 | 0.09720 | 0.146103 | H2 | 0.04730 | 0.076123 |
| | | | O2’ | -0.61390 | -0.626760 | N3 | -0.69970 | -0.744123 |
| | | | HO’2 | 0.41860 | 0.462358 | C4 | 0.30530 | N.A. |
| | | | O3’ | -0.65410 | -0.65410 | N4 | N.A. | 0.603011 |
| | | | H3T | 0.43760 | 0.437600 | | | |

#### Selecting Reference Structures

The reference states used for the reaction coordinate or the implementation of TMD need to be close to equilibrium but not at equilibrium, since we want to sample both sides of equilibrium region along the RC, while the reference structures correspond to the two ends of the RC$(-RC_{max}, +RC_{max})$, with $RC_{max} = \delta RMSD(X_{Openref}, XClosedref)$. The reference structures or states are selected using the first 50ns of the unrestrained NPT simulations, and the correspondingly defined RCs for the open and closed equilibrated structures need to satisfy the conditions below:

$$\begin{aligned}
\delta RMSD(X_{\text{Open equi}}) =& RMSD(X_{\text{Open equi}}, X_{\text{Open ref}}) - \\
& RMSD(X_{\text{Open equi}}, X_{\text{Closed ref}}) \\
\delta RMSD(r_{\text{Closed equi}}) =& RMSD(X_{\text{Closed equi}}, X_{\text{Open ref}}) - \\
& RMSD(X_{\text{Closed equi}}, X_{\text{Closed ref}})
\end{aligned} \tag{1}$$

Where the requirement is:

$$
\begin{aligned}
-RC_{max} < RC(X_{\text{Open equi}}) < 0 \\
0 < RC(X_{\text{Closed equi}}) < +RC_{max}
\end{aligned}
\tag{2}
$$

Where the RC is $\delta RMSD$ specified in equation (1).

#### TMD + Umbrella Sampling Parameters

Table S2: Summary of target MD and Umbrella Sampling parameters. The force constant used from the TMD simulations were carried over and use for the respective umbrella sampling simulations. The largest force constant was used for the ATP simulations and smallest for RDV (stacking). Where the () in RDV indicate the initial binding structure (open for the active site open state).

| RC | Force Constant $\left(\frac{kcal}{mol\mathring{A}^2}\right)$ | RC Range(Å) | Number of Windows |
|---|---|---|---|
| Motifs + ATP | 501 | -1.2 to 1.3 | 27 |
| Motifs + ATP + Template | 501 | -1.1 to 1.3 | 26 |
| Motifs + RTP(Open Stacking) | 125 | -1.0 to 1.0 | 21 |
| Motifs + RTP(Open Stacking) + Template | 125 | -1.6 to 1.6 | 34 |
| Motifs + RTP(Open Base-pairing) + Template | 250 | -1.5 to 1.5 | 32 |

#### APBS (Adaptive Poisson-Boltzmann Solver)

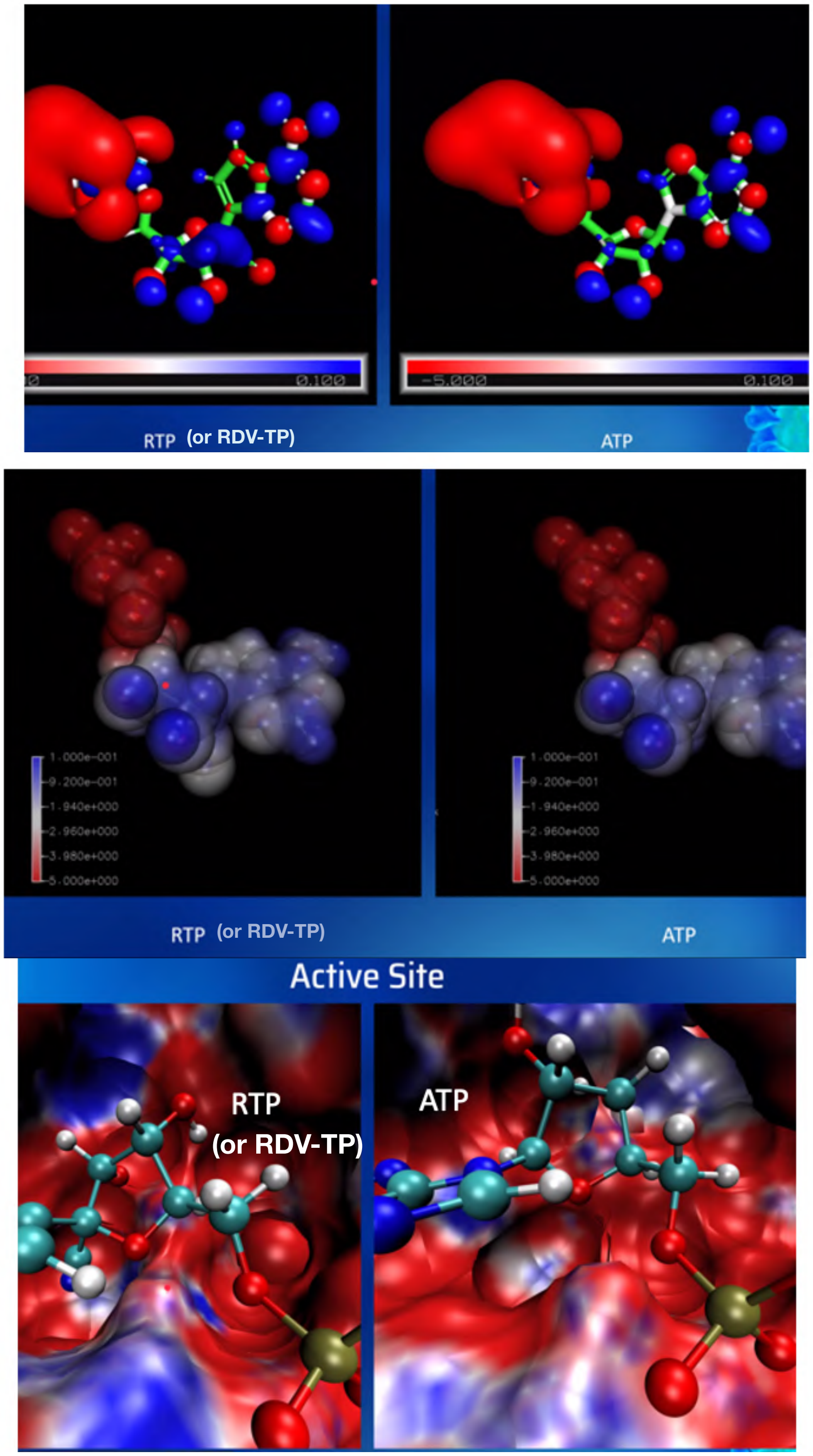


Figure S2: The electrostatic potential generated by solving the Poisson-Boltzmann equation using the APBS solver [7]. The potential due to partial charges assigned for RDV-TP (or RTP) in comparison with that of ATP (left). The potentials around the active site of the CoV-2 RdRp (PDB: 7BV2) with RDV-TP and ATP inserted, respectively.

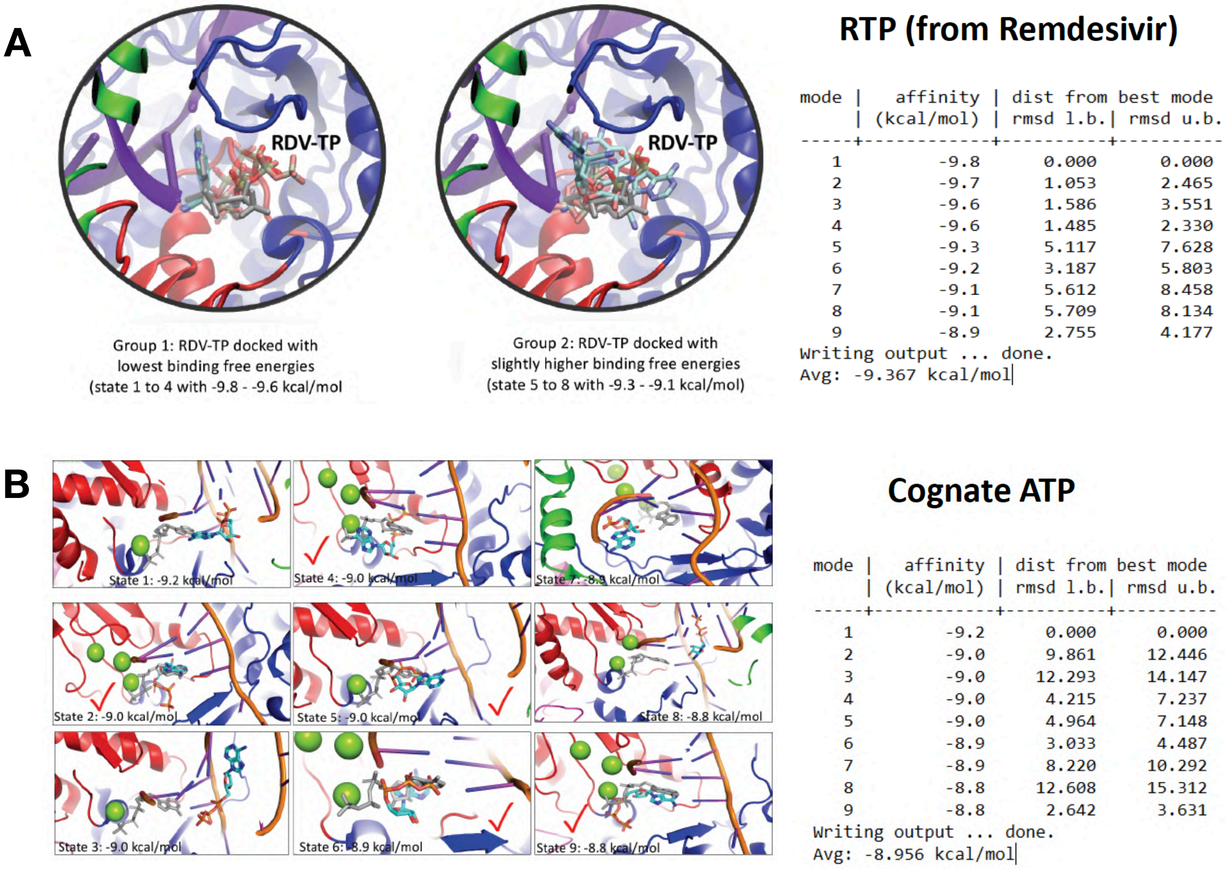


Figure S3: Docking of RDV-TP and ATP onto a modeled initial binding (active-site open) structure of SARS-CoV-2 RdRp (PDB: 7BTF). **A.** RDV-TP docking show comparatively stabilized docking structures (grouped into two). The palm, fingers, thumb subdomains are shown in red, blue, and green, and RNA in violet. The modeled RDV-TP (positioning from PDB: 7BV2) is shown in gray. **B.** ATP docking shows diverse configurations and less stabilized configurations. The obtained docking energetics are listed on the right side (using AutoDock [2]

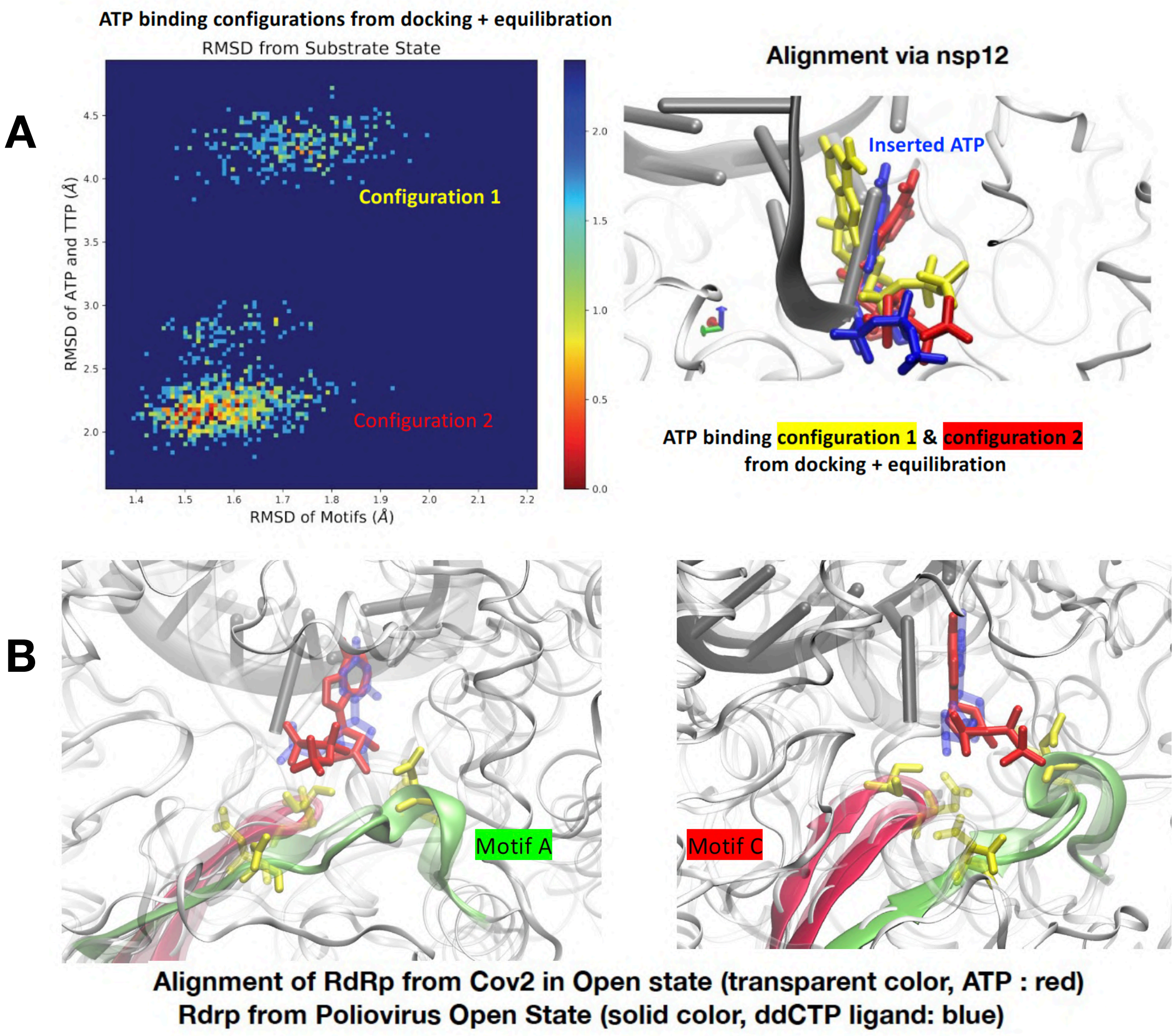


Figure S4: Examining ATP binding configurations to the open form active site CoV-2 RdRp. **A.** The MD equilibration of the optimal docking complex of ATP to the CoV-2 RdRp structure. The equilibrated configurations were measured by RMSDs for both structural motifs (A-G) and ATP+template TTP with respect to the substrate insertion complex. Two dominant configurations of ATP have been identified, both of which are quite close to the insertion configuration. **B.** The alignment of the ATP bound open form CoV-2 RdRp with that of the poliovirus (PV) RdRp, shown in two angles for better view. We further conducted equilibrium MD simulations on the optimized docking complex of ATP. The results show that even upon the docking and equilibration, the stabilized configurations of ATP still converge to be very close to the initial modeled ones (from the insertion structural complexes; see Fig S6A). Interestingly, even we chose another reference structure in docking (e.g. using the pre-insertion structure of T7 RNAP [8]), we could still obtain a docking configuration overlapping well with the insertion ATP. Hence, it justifies that the above constructed ATP and RTP initial binding or the active-site open RdRp structural complexes are reasonable. A further comparison show that ATP binding configuration in our constructed open form RdRp complex is similar to that being captured in the PV RdRp.

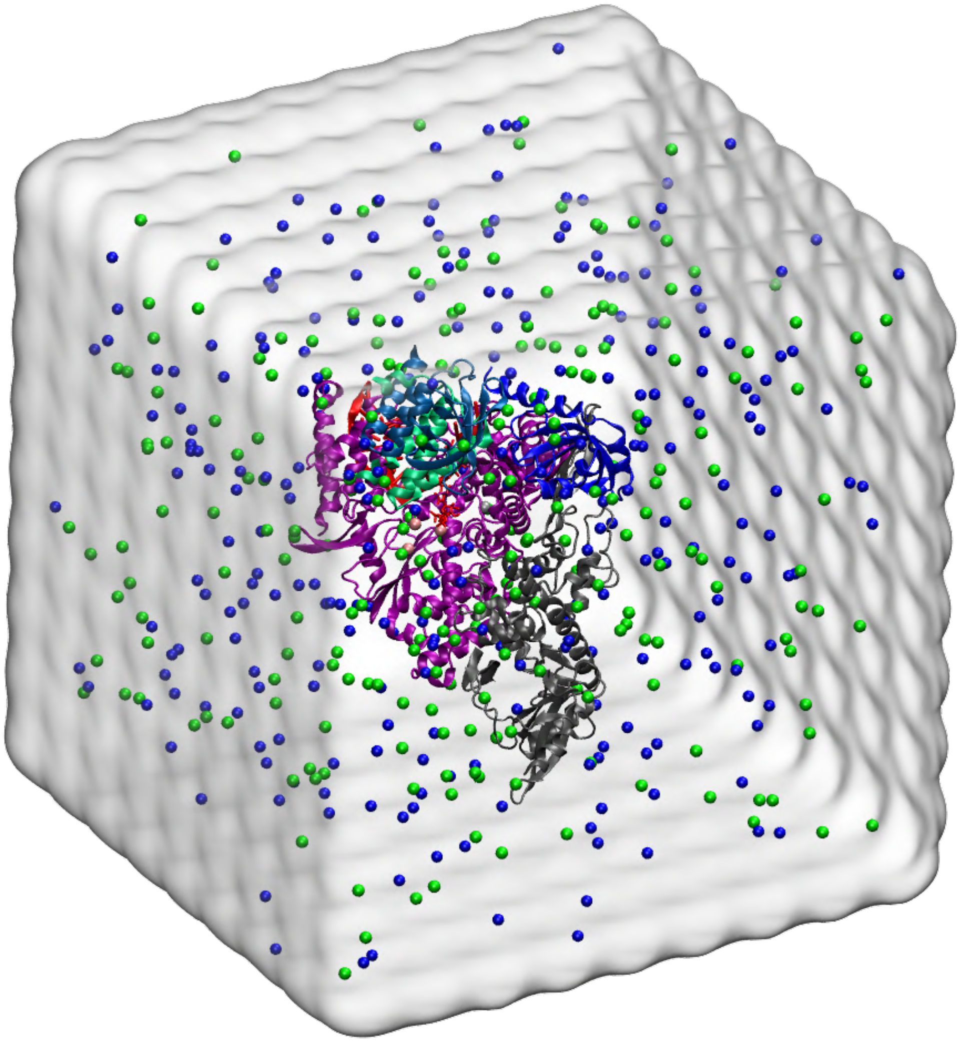

Figure S5: All-atom molecular dynamics simulation box. The size of the avg. box: 15.7nm x 15.7 nm x 15.7 nm, containing an avg. of: 382,000 atoms atoms (118,484 water molecules, 266 sodium and 230 chloride ions).

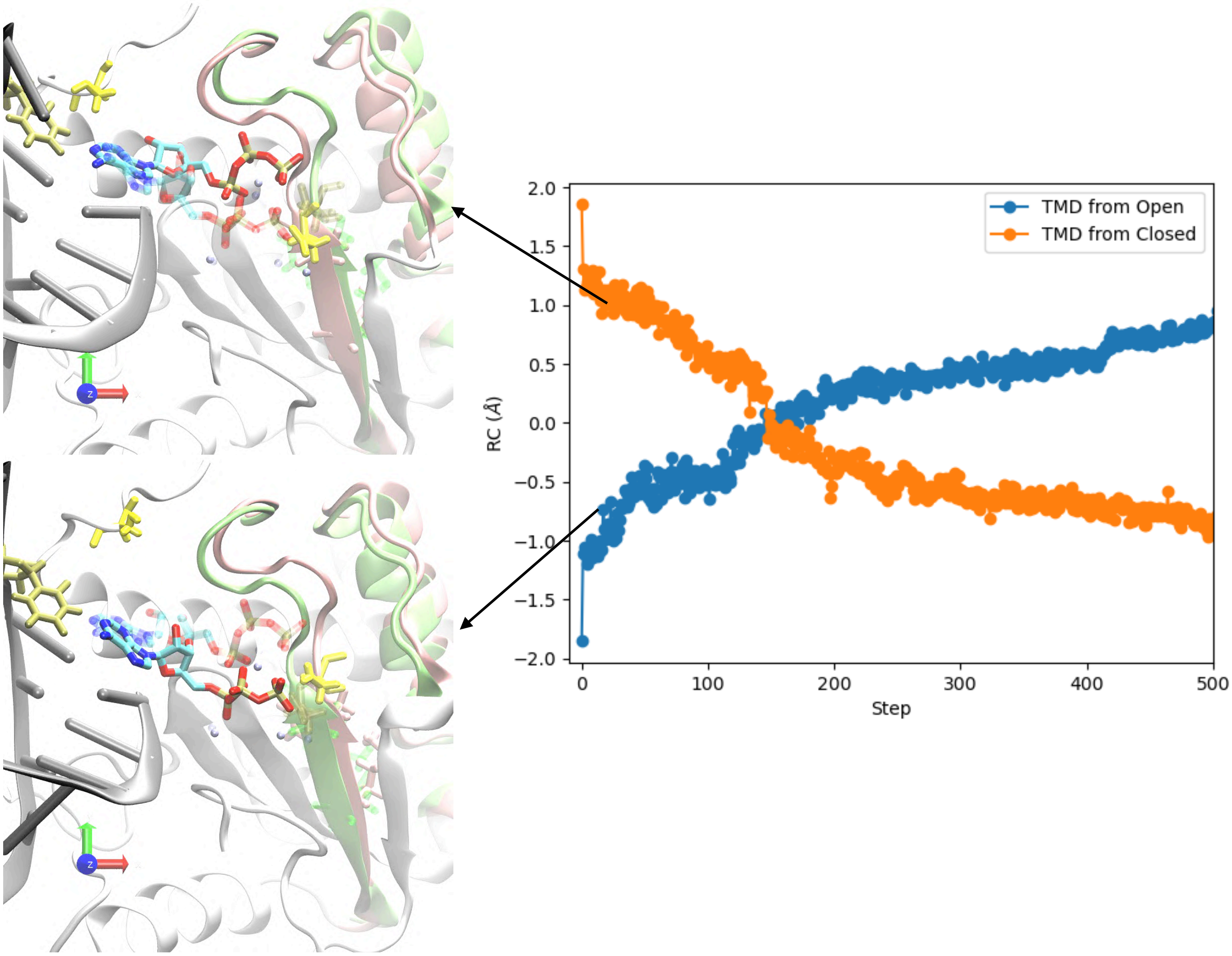


Figure S6: Implementation of targeted molecular dynamics (TMD) simulations for constructing NTP insertion path to be utilized in the umbrella sampling simulations. (Left) The initial and final structures of each respective paths: backward (bottom image) and forward (upper image). With motifs A/D colored pink for the starting structure and green for the target structure. Representations colored to compare with PV RdRp in [9]. (Right) The implementation of the TMD simulations forward and backward. Where TMD from open refers to beginning TMD from the initial binding complex (forward path) and TMD from closed is beginning TMD from insertion complex (backward path). Structures are selected every 0.1 Å from the two paths until their intersection for the umbrella sampling simulation windows.

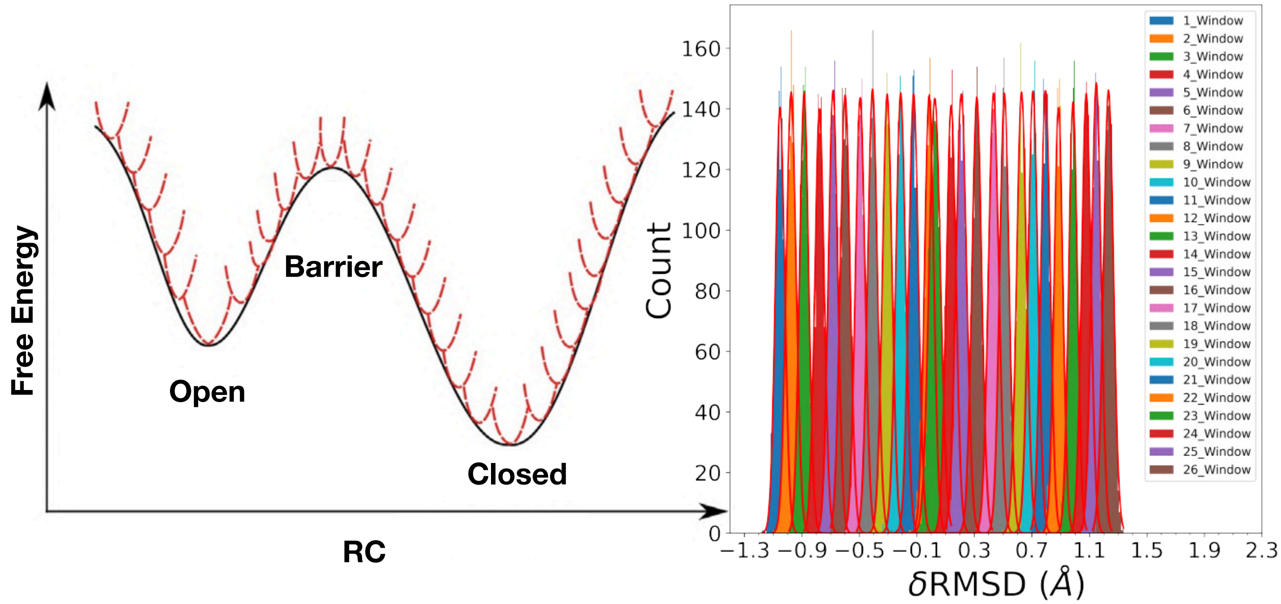


Figure S7: Conducting the umbrella sampling simulations for NTP insertion. (Left) The schematics of conducting the umbrella sampling simulation strategies (figure adapted from [10]). (Right) The overlap of simulated windows, where the RC is centered every 0.1Å with the initial simulation structure taken from the forward/backward TMD paths see **Fig S6**.

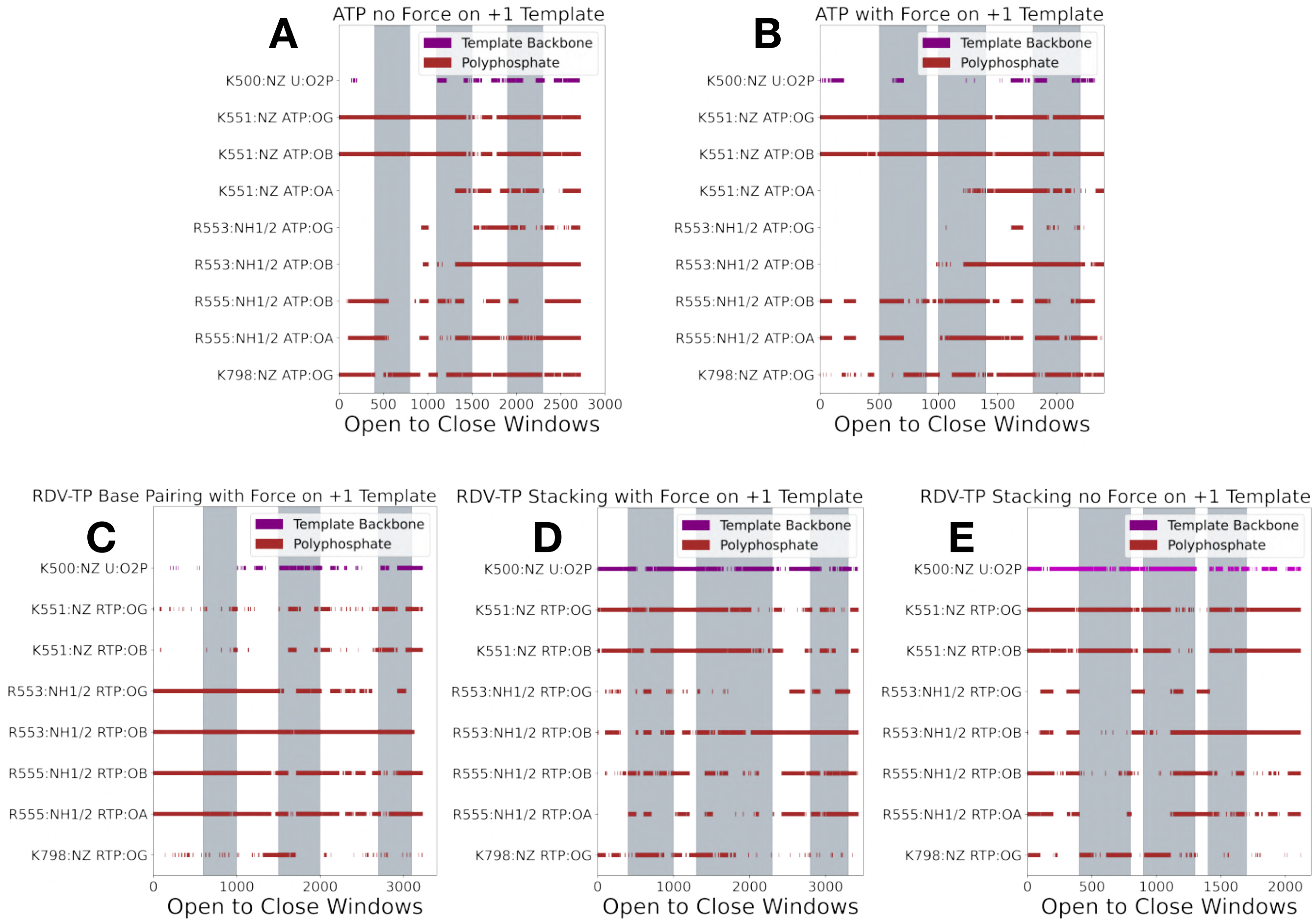


Figure S8: From the hydrogen bond analysis (see main text **Fig 3-7**) we can identify the residues (Lys and Arg) which can form salt bridges along the polyphosphate. Further exploration identified no other salt bridges along the polyphosphate of ATP/RDV-TP or the +1 template nt. Measurements are conducted as discussed in the hydrogen bond analysis but with a cutoff of 5Å to identify a salt bridge. Measurements are done from the side chain heavy atoms from Lys/Arg and oxygen atoms along the polyphosphate.

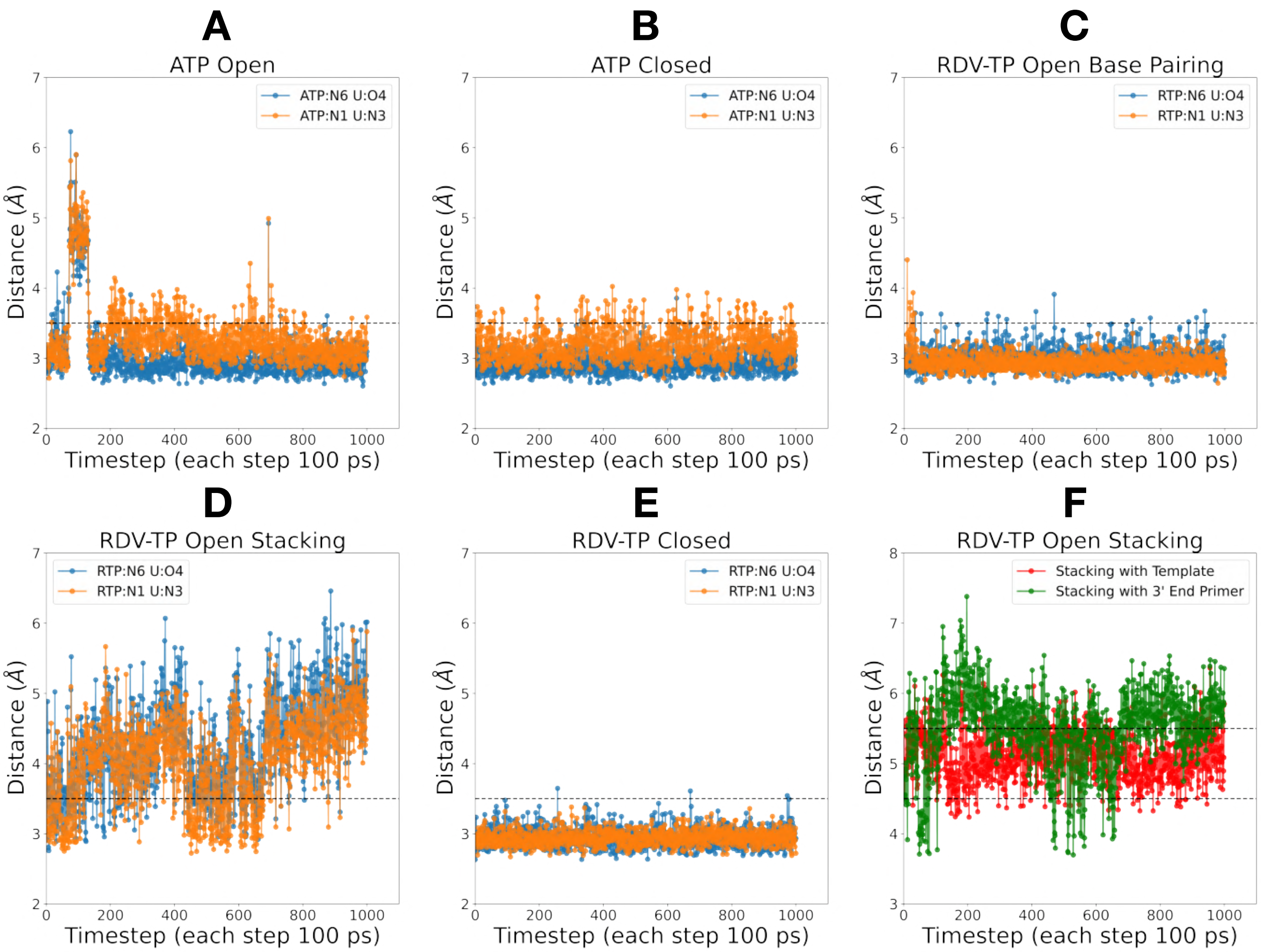


Figure S9: Expected hydrogen bond (HB) distance between ATP/RDV-TP and the +1 template Uracil (**A-E**.) The distance pairs measured are the heavy atoms from the nucleotide triphosphate N6 with U:O4 and N1 and U:N3. The dashed black line indicate the estimated cutoff (3.5 Å) for a HB. The insertion complexes (**B&E**) form significantly more stable HB. An RDV-TP initial-binding forms a stacking configuration (**D**) in which hydrogen bonds are rarely formed. **F** Stacking is measured from this trajectory by taking the center of mass of the 6-numbered rings on the bases (RDV and Uracil for +template / 3' end primer). Reasonable stacking is observed between ∼ 4.5Å - 5.5Å

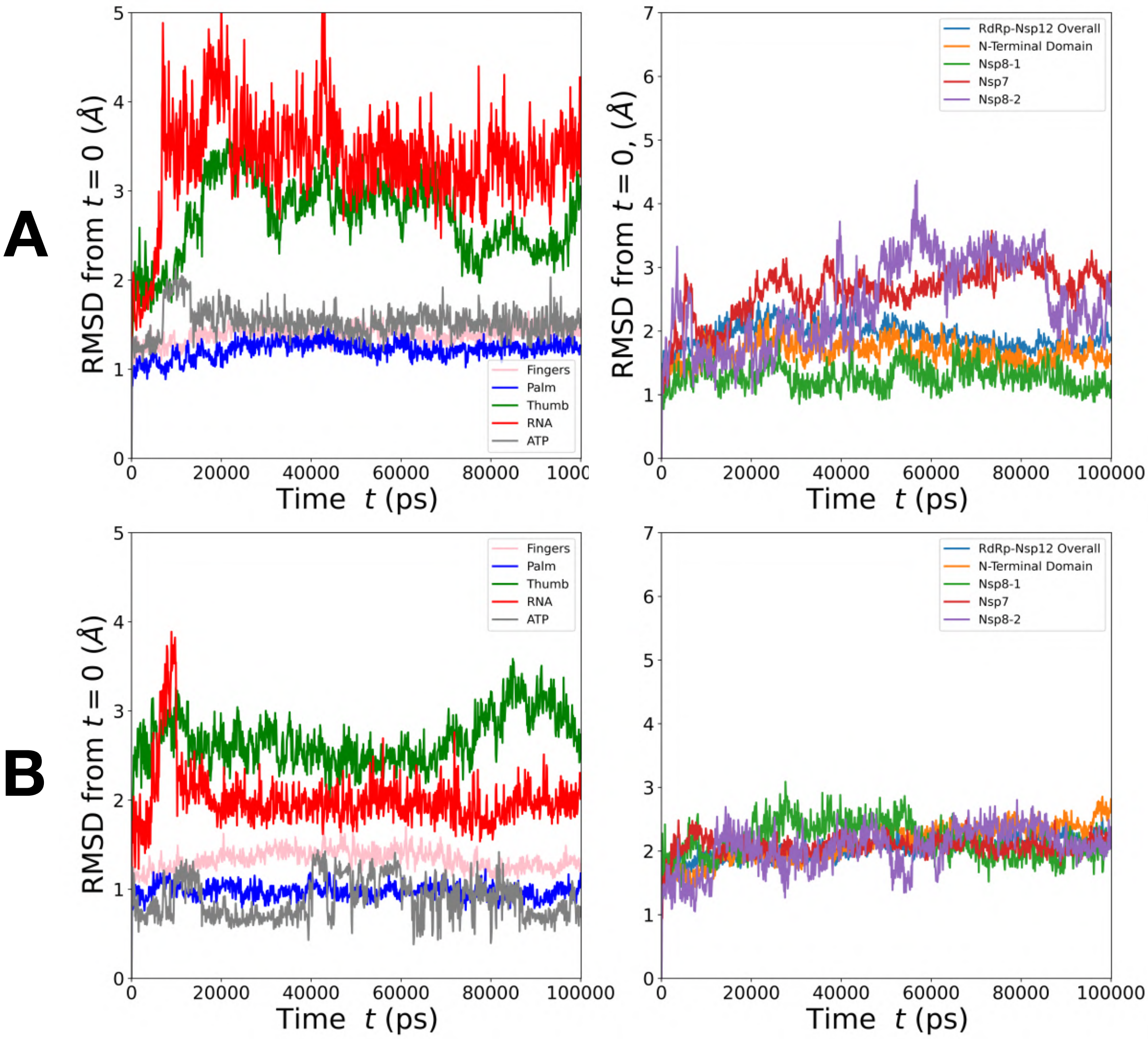


Figure S10: Measured RMSD from equilibration simulations. (Left) Subdomains, RNA, and ATP. (Right) Cofactors, Nsp12 and N-Terminus domain. **A** ATP initial binding complex. **B** ATP insertion complex. The insertion complex is more stable than the initial binding complex

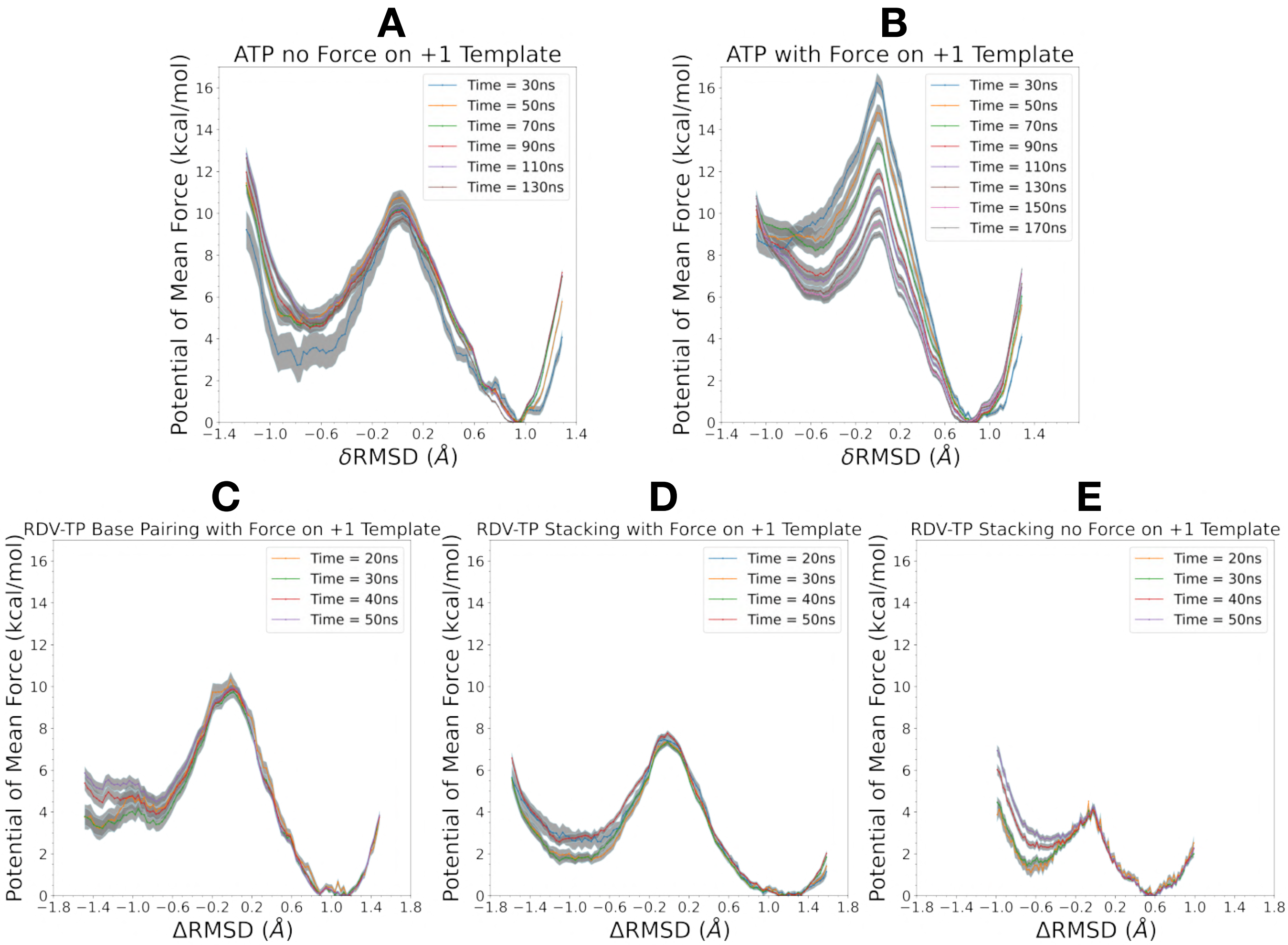


Figure S11: Convergence plots of all PMFs constructed with bootstrapping error analysis for each set of data. As more data is accumulated with the extended simulations the PMF construction, the PMF further converges and estimated error decreases. Early data collected is removed for pre-equilibration, 20ns for ATP and 10ns for RDV-TP systems. **A&B** ATP PMF's with no force on template and with force on template, respectively. **C** RDV-TP base pairing with force on +1 template nt. **D** RDV-TP stacking with force on + 1 template nt. **E** RDV-TP stacking with no force on + 1 template. **D-E** Only 50ns of data from each window was needed to reach convergence.

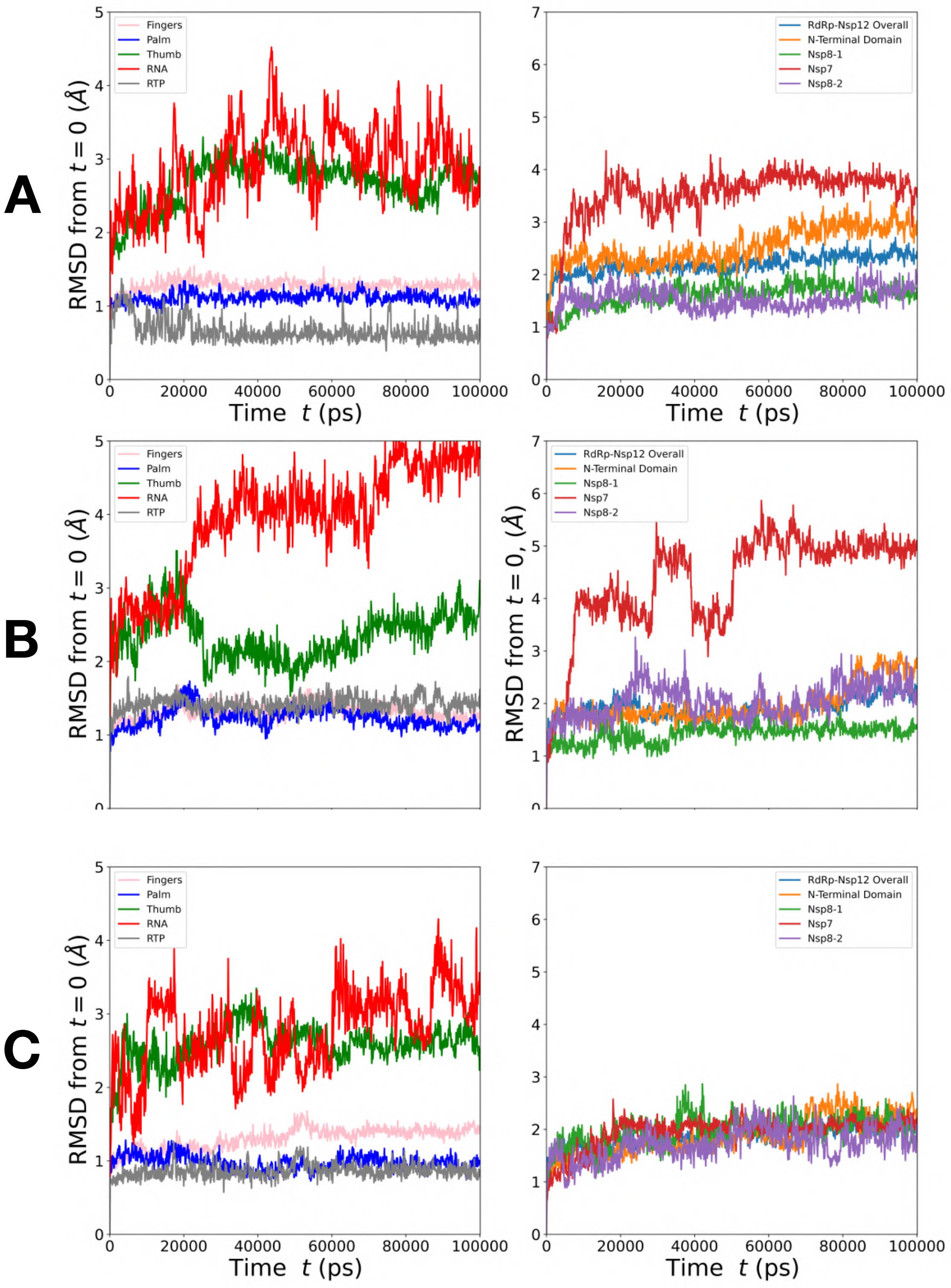


Figure S12: Measured RMSD from RTP (RDV-TP) equilibration simulations. (Left) Subdomains, RNA, and NTP. (Right) Cofactors, Nsp12 and N-Terminus domain. **A** RDV-TP initial binding base pairing configuration. **B** RDV-TP initial binding stacking configuration. **C** RDV-TP insertion complex. Similar to ATP insertion complex **Fig S10B** the RDV-TP closed complex is stable than the both of its open configurations.

**Movie S1**

Forward path TMD for ATP, the target structure is the ATP inserted. The representation follows the same described in **Fig S6** in reference to PV [9].

**Movie S2**

Forward path TMD for RDV-TP (initial binding stacking and no force on template nt), the target structure is the RDV-TP inserted. The representation follows the same described in **Fig S6** in reference to PV [9].

[5] Junmei Wang, Piotr Cieplak, and Peter A Kollman. How Well Does a Restrained Electrostatic Potential (RESP) Model Perform in Calculating Conformational Energies

of Organic and Biological Molecules? Keywords: additive force field; nonadditive force field; restrained electrostatic potential (RESP); torsion. Technical Report 12, 2000.

[6] Wendy D Cornell, Piotr Cieplak, / Christopher, I Bayly, Ian R Gould, Kenneth M Merz, David M Ferguson, David C Spellmeyer, Thomas Fox, James W Caldwell, and Peter A Kollman. A Second Generation Force Field for the Simulation of Proteins, Nucleic Acids, and Organic Molecules. Technical report, 1995.

[7] Elizabeth Jurrus, Dave Engel, Keith Star, Kyle Monson, Juan Brandi, Lisa E. Felberg, David H. Brookes, Leighton Wilson, Jiahui Chen, Karina Liles, Minju Chun, Peter Li, David W. Gohara, Todd Dolinsky, Robert Konecny, David R. Koes, Jens Erik Nielsen, Teresa Head-Gordon, Weihua Geng, Robert Krasny, Guo Wei Wei, Michael J. Holst, J. Andrew McCammon, and Nathan A. Baker. Improvements to the APBS biomolecular solvation software suite. *Protein Science*, 27(1):112–128, 1 2018.

[8] Dmitry Temiakov, Vsevolod Patlan, Michael Anikin, William T. McAllister, Shigeyuki Yokoyama, and Dmitry G. Vassylyev. Structural basis for substrate selection by T7 RNA polymerase. *Cell*, 116(3):381–391, 2 2004.

[9] Grace Campagnola, Seth McDonald, Stéphanie Beaucourt, Marco Vignuzzi, and Olve B Peersen. Structure-Function Relationships Underlying the Replication Fidelity of Viral RNA-Dependent RNA Polymerases. *Journal of Virology*, 89(1):275–286, 2015.

[10] Qinghua Liao. Enhanced sampling and free energy calculations for protein simulations. In *Progress in Molecular Biology and Translational Science*, volume 170, pages 177–213. Elsevier B.V., 1 2020.
